# Host cell membrane capture by the SARS CoV-2 spike protein fusion intermediate

**DOI:** 10.1101/2021.04.09.439051

**Authors:** Rui Su, Jin Zeng, Ben O’Shaughnessy

## Abstract

Cell entry by SARS-CoV-2 is accomplished by the S2 subunit of the spike S protein on the virion surface by capture of the host cell membrane and fusion with the viral envelope. Capture and fusion require the prefusion S2 to transit to its potent, fusogenic form, the fusion intermediate (FI). However, the FI structure is unknown, detailed computational models of the FI are unavailable, and the mechanisms and timing of membrane capture and fusion are not established. Here, we constructed a full-length model of the CoV-2 FI by extrapolating from known CoV-2 pre- and postfusion structures. In atomistic and coarse-grained molecular dynamics simulations the FI was remarkably flexible and executed large bending and extensional fluctuations due to three hinges in the C-terminal base. Simulations suggested a host cell membrane capture time of ∼ 2 ms. Isolated fusion peptide simulations identified an N-terminal helix that directed and maintained binding to the membrane but grossly underestimated the binding time, showing that the fusion peptide environment is radically altered when attached to its host fusion protein. The large configurational fluctuations of the FI generated a substantial exploration volume that aided capture of the target membrane, and may set the waiting time for fluctuation-triggered refolding of the FI that draws the viral envelope and host cell membrane together for fusion. These results describe the FI as a machinery designed for efficient membrane capture and suggest novel potential drug targets.

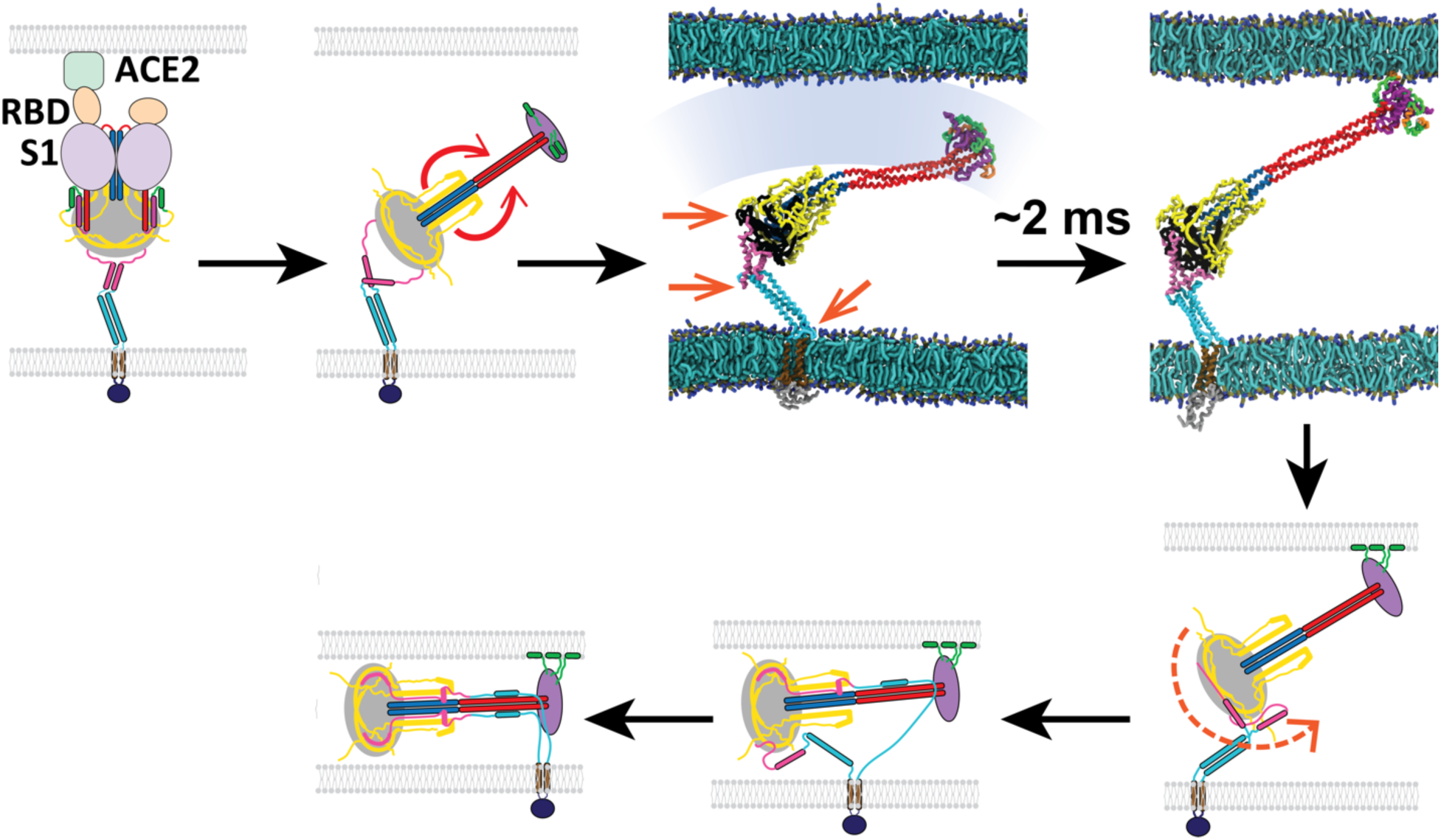

## Introduction

The COVID-19 global pandemic is this century’s third coronavirus epidemic following SARS-CoV in 2002 and Middle East respiratory syndrome coronavirus (MERS-CoV) in 2012, suggesting that coronaviruses will remain a global health threat for the foreseeable future. The responsible pathogen is the severe acute respiratory syndrome coronavirus 2 (SARS-CoV-2), a ∼100-nm diameter betacoronavirus^1^ whose lipid envelope encloses a positive-sense single-stranded RNA genome complexed with the nucleocapsid (N) protein. The lipid envelope houses the spike (S) glycoprotein, the envelope (E) protein and the membrane (M) protein^2^. The S protein is a trimeric class I fusion protein that catalyzes entry into lung, nasal mucosa or small intestine cells and has two subunits, S1 and S2^3, 4^. Following S1-mediated binding to host cell membrane Angiotensin-Converting Enzyme 2 (ACE2) receptors and proteolytic activation at locus S2′ by TMPRSS2 on the plasma membrane (PM)^5^ or by endosomal cathepsins B and L^6^, S1 and S2 are thought to dissociate, releasing S2.

Entry is the job of the S2 subunit, by fusion of the viral envelope and host cell membrane. To become fusion competent, S2 must first undergo a major structural transition from its prefusion state to the potent, fusogenic form, the extended fusion intermediate (FI)^7, 8^ which bears three N-terminal fusion peptides (FPs) that capture the host cell membrane. Subsequent refolding of the fusion intermediate into its postfusion configuration^9^ pulls the viral envelope and target membrane together for fusion and delivery of viral genomic material^10^.

The fusion intermediate is the unsheathed weapon of CoV-2 entry, but the mechanism and timescales of FI-mediated host cell membrane capture are unknown. Little is known about this critical machinery. While the prefusion and postfusion CoV-2 S protein structures are known from cryo-EM and x-ray crystallography^3, 9, 11, 12^, the FI structure is undetermined. Extended intermediate states of class I fusion proteins have generally proved experimentally elusive, likely due to their estimated sec-min lifetimes, far shorter than the pre- or postfusion lifetimes^7, 13, 14^ . Indeed, no class I fusion protein intermediate had been visualized until very recently, when the HIV-1 gp41 intermediate, the parainfluenza F intermediate and the influenza HA2 intermediate^14–16^ were visualized with cyro-EM for the first time.

The spike protein is a target for vaccines, therapeutic antibodies and other antivirals. Most current recombinant neutralizing or vaccine-elicited antibodies to CoV-2 bind S1^17–20^. Viral fusion inhibitors targeting S2^12, 21^ and the fusion-executing subunits of other class I fusion proteins have been developed, including one FDA-approved drug against HIV^22–24^. Importantly, recently emerged CoV-2 variants harboring spike protein mutations, B.1.351 (South African variant) and P.1 (Brazilian variant), escaped from two S1-targeting antibodies one of which has FDA emergency authorization, whereas the efficiency of S2-targeting antivirals was unaffected^25^. Moreover, S2 is conserved among coronaviruses^26^. Thus, unveiling the mechanisms of the CoV-2 S2 fusion intermediate will be vital in the search for robust and pan-coronavirus antiviral drugs.

Detailed computational studies of the CoV-2 FI and the kinetics of FI-mediated membrane capture are unavailable, to our knowledge. However the prefusion S protein was atomistically simulated^27–29^, one study revealing a highly flexible prefusion structure due to three hinge-like regions, consistent with cryo-ET^28^. Membrane binding by the isolated FP was simulated^30–32^, including Ca^2+^-dependent binding which involved the conserved coronavirus LLF motif in the N-terminal FP helix^30^. A coarse-grained model of the whole virion was developed^33^. For other class I fusion proteins, a model structure of the Ebola GP2 extended intermediate was constructed^34^ and local transitions of the influenza HA and HIV gp41 intermediates were modelled^35, 36^.

Here, we constructed a full-length structural model of the CoV-2 FI by extrapolating from known CoV-2 pre- and postfusion structures. The model suggests a “loaded spring” mechanism triggers the prefusion-to-FI transition, similar to that for influenza HA thought triggered by folding of the B-loop into a helix^37^. We studied membrane capture by the FI, using all-atom (AA) and MARTINI coarse-grained (CG) molecular dynamics (MD) simulations to access timescales up to a ms. Simulations showed the FI is highly flexible, subject to large orientational and extensional fluctuations due to three hinges in the C-terminal base closely related to the prefusion hinges^28^. These fluctuations greatly increased the volume swept out by the FI, helping it capture target cell membrane. A critical N-terminal amphiphilic helix in the FP mediates membrane binding, but we find FP-only simulations severely mispresent the kinetics, as membrane capture is far slower in the native structure with the FP attached to the FI. Our CG simulations suggest FI-mediated membrane capture requires ∼ 2 ms, a critical step on the pathway to fusion. In addition to facilitating membrane capture, we propose that large FI fluctuations set the timing of fluctuation-triggered FI refolding on the pathway to membrane fusion. Our work identifies several novel potential drug targets.

## Results

### Model of the CoV-2 spike protein fusion intermediate

Entry of SARS-CoV-2 is catalyzed by the trimeric S protein, consisting of three S1 heads sitting upon a trimeric S2 stem^3^. Following binding of S1 to the host cell ACE2 receptor^5^, S1 dissociates from S2^38^ and the prefusion S2 trimer undergoes a major structural transition to its potent, fusogenic form, the fusion intermediate (FI)^7^ (Figure 1a).

**Figure 1.**
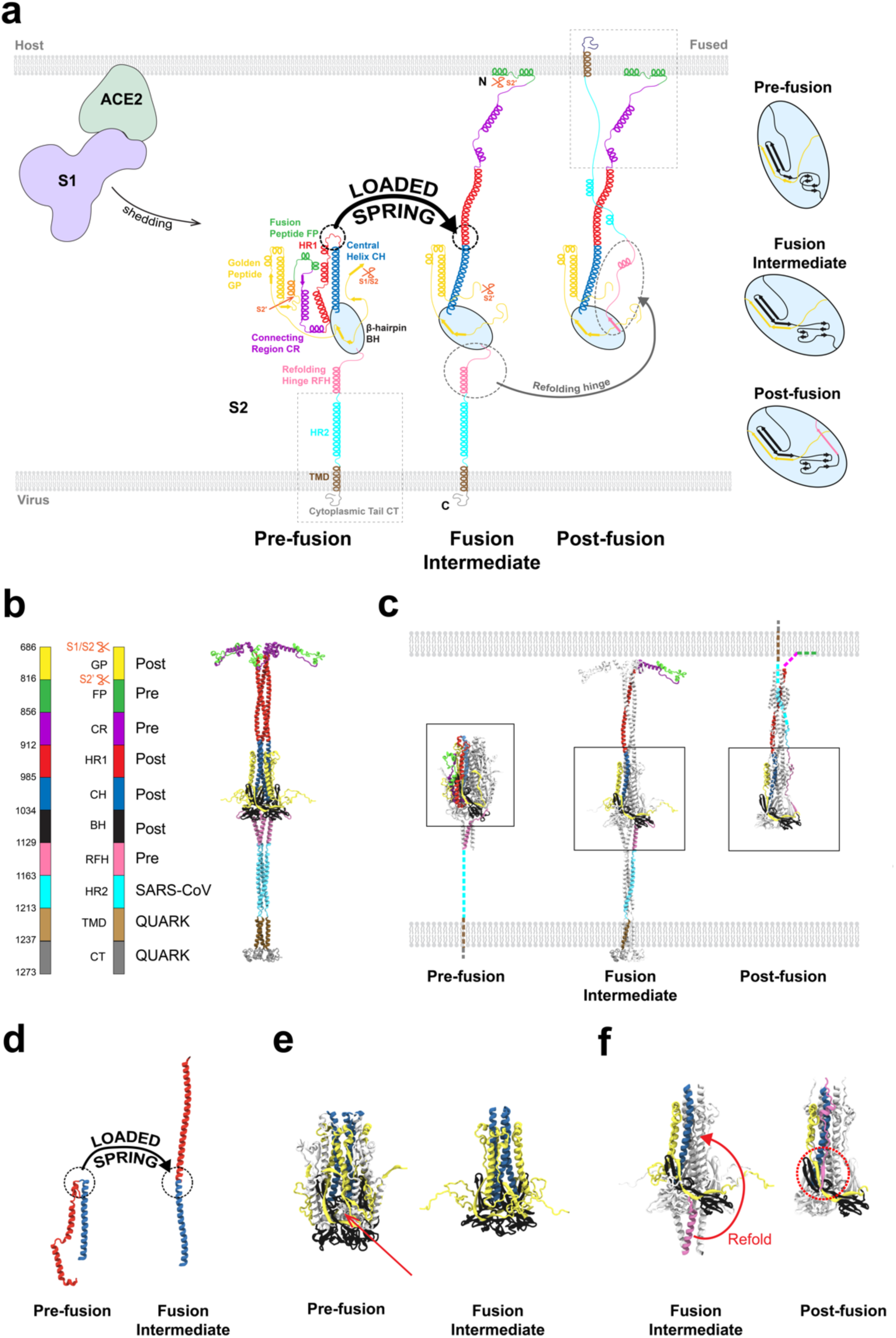
Model of the SARS-CoV-2 spike protein fusion intermediate. **(a)** Model of the fusion intermediate (FI) of the CoV-2 spike (S) protein, schematic. One protomer of the S trimer is shown. Schematics of the pre- and postfusion S2 states from the known structures, other than unknown regions (boxed). Details of BH and adjacent domains, at right. Following dissociation of S2 from S1, the unstructured pre-fusion HR1 loops become helical (loaded spring release), giving the HR1-CH backbone that thrusts the FPs toward the host cell membrane. The FI subsequently refolds into the postfusion structure, driven by structural changes in RFH and chaperoned by GP, with RFH and HR2 providing leashes that pack the HR1-CH backbone grooves. **(b)** Model of the FI of the CoV-2 spike (S) protein, exact structure. Source structures for each S2 subunit domain are indicated, either CoV-2 prefusion (PDB: 6XR8), CoV-2 postfusion (PDB: 6XRA) or HR2 of CoV (PDB: 2FXP). Transmembrane domain (TMD) and cytoplasmic tail (CT) structures predicted by QUARK. **(c)** Comparison between predicted FI structure and known crystal structures of the prefusion (PDB: 6XR8) and postfusion (PDB: 6XRA) CoV-2 S2 subunit. One protomer highlighted in color. Dashed lines: missing domains from partially solved crystal structures. **(d)** Details of loaded spring transition. **(e)** β-hairpin (BH) domains in the known prefusion and predicted FI structures (boxed regions of (c)). The RFH domain is omitted from the FI BH for clarity. Following the loaded spring transition, HR1, CR and FP (shown faint in the prefusion BH) vacate their prefusion locations in BH. The resultant cavity (arrow) would presumably be unstable. We assume the FI adopts the more compact postfusion BH structure (right). **(f)** The golden peptide (GP) domain chaperones refolding of the fusion intermediate (FI) into the postfusion structure. Blowups of boxed regions in (c) are shown. Refolding of the refolding hinge (RFH) domain is guided by GP. RFH forms a parallel β-strand with GP (red circle), the RFH unstructured portion packs the CH-GP groove, and RFH helices interact with two GP helices. Colored BH and CH belong to one protomer; colored RFH belongs to a different protomer.

We predicted the structure of the FI based on information from the partially solved pre-and postfusion cryo-EM structures^9^ (Figures 1b,c). A natural question is whether formation of the FI from its prefusion state uses a load spring mechanism similar to that used by influenza HA for the analogous transition^37^. We hypothesized that in S2 a transition converts the prefusion heptad repeat 1 (HR1) domain into a continuous alpha helix, itself continuous with the downstream central helix (CH). That is, all unstructured loop domains in HR1 become helical and rotate like torsional springs, straightening HR1 (Figures 1a,d). The result is a three-helix HR1-CH coiled coil in the FI trimer, the mechanical backbone of the FI. The hypothesis is tantamount to assuming CH and HR1 adopt their postfusion structures in the FI, since the postfusion HR1 and CH are continuous helices in a trimeric coiled coil^9^ (Figure 1c).

The S2′ site is assumed cleaved before this transition, exposing the FP N-terminus ready for host membrane capture, consistent with CoV-2 lung cell entry being blocked by inhibition of the serine protease TMPRSS2 that cleaves S2′ at the host cell plasma membrane^5^. This cleavage disconnects the golden peptide (GP) domain, but GP remains physically attached to S2^9^. (We tested an uncleaved model with connected GP and FP, but the FPs were sequestered and unable to access the target membrane, Figure S1.)

We assumed the β-hairpin (BH) domain adopts its postfusion configuration in the FI. Straightening of HR1-CH pulls these domains away from their prefusion locations and would leave large destabilizing cavities in BH, favoring transition to the postfusion configuration where BH is raised to fill the cavities and assembles into a pyramidal base (Figure 1e). The same is assumed of GP, since a GP β-strand interacts strongly with an antiparallel β-sheet in BH (Figure 1a). GP also contributes two small helices, and a long helix in a CH coiled coil groove completing a six-helix bundle (Figures 1a, e) providing structural support at the base of the CH-HR1 backbone (see below).

Downstream of BH the refolding hinge (RFH) domain, remote from the loaded springs, was taken as the prefusion structure, while we used the HR2 structure of SARS-CoV from NMR^39^ whose HR2 sequence is the same as that of CoV-2^12^. The unknown TM and CT structures were predicted by QUARK^40^.

These components were integrated into the predicted CoV-2 FI structure shown in Figure 1b (see Methods). The model implies that subsequent refolding of the FI to the postfusion conformation occurs by the unstructured N-terminal loop of the refolding hinge (RFH) domain folding into BH by contributing a β-strand to an antiparallel β-sheet (Figures 1a, f). The remainder of RFH folds back as a leash packing a GP-CH groove in the six-helix bundle, ending in a small helix that attaches between the two small GP helices of the other two protomers, oriented almost perpendicular to the HR1-CH backbone. Refolding is completed when the helical HR2 becomes partially unstructured to pack a second leash into a HR1-CH groove and supply one helix to a six-helix postfusion bundle with HR1, the fusion core^9^.

### All-atom simulation of the fusion intermediate

Using complementary atomistic and coarse-grained MD methods, we tested the FI model of Figure 1b and measured its configurational statistics and dynamics (see Methods).

During ∼ 0.4 µs of AA simulation the basic secondary structure remained unaltered (Figure S2), lending credibility to the model. Far from the rigid extended object one might anticipate given its long helical domains (Figure 1b), the FI was highly flexible and underwent large configurational fluctuations, adopting bent configurations without structural damage (Figure 2). The structural robustness was due to energy-absorbing features. The RFH-HR2 base region downstream of the BH domain was highly flexible, allowing large tilt (Figure 2b and Supplementary Movie 1). Relative to the prefusion structure the three RFH helices, known as the stem helices, became splayed with separated N-termini, in an inverted tripod suspension system that buffered large displacements of the upstream BH and backbone. The HR2 helices became partially unstructured, a structural plasticity that helped the FI tilt to greater angles at the membrane (Figure 2b).

**Figure 2.**
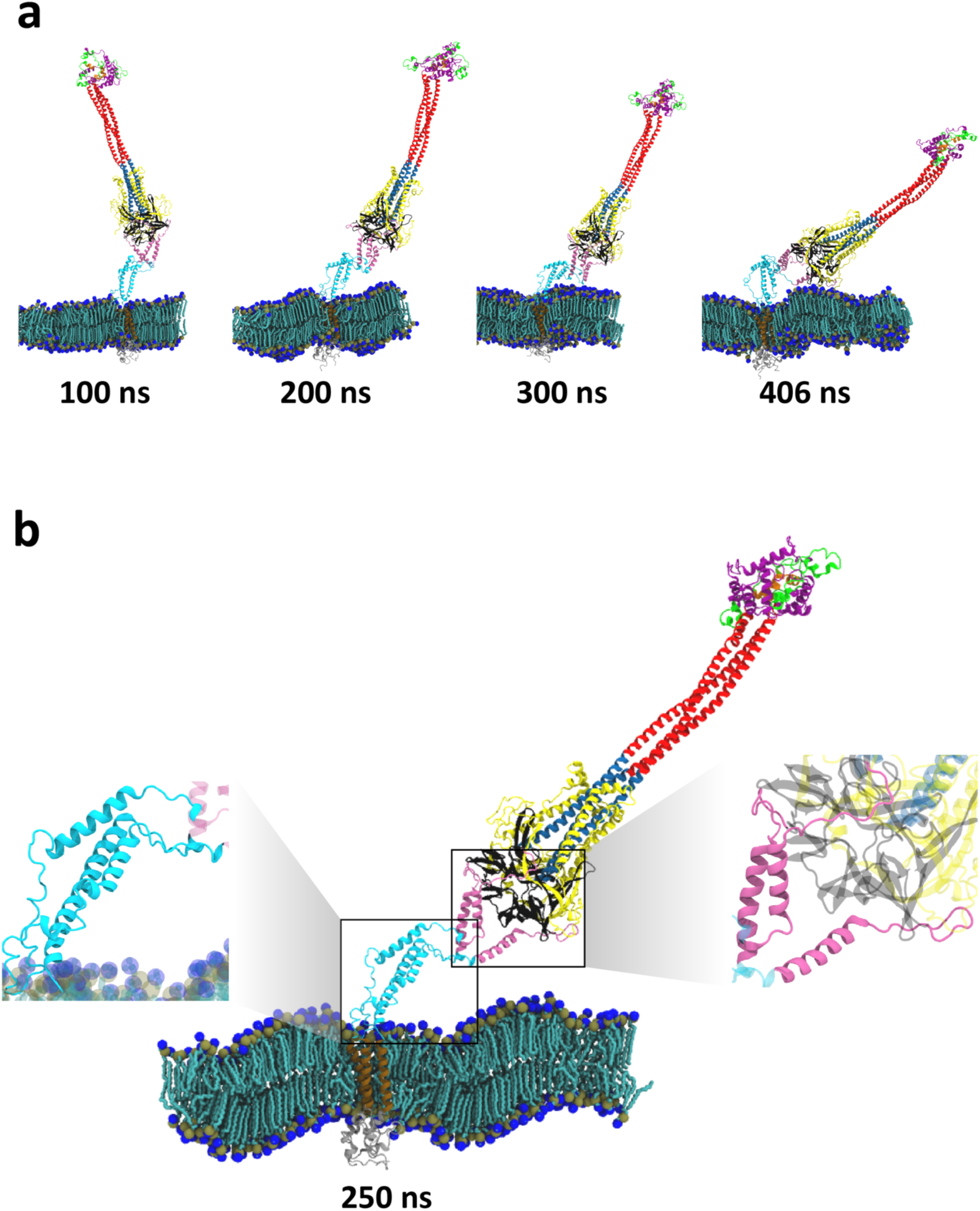
All-atom simulation of the SARS-CoV-2 fusion intermediate. Color code for this and all subsequent figures, as for Figure 1. In addition, the N-terminal helices of the fusion peptides are shown orange. **(a)** Snapshots of the FI during the ∼ 0.4 µs all-atom simulation of the model of Figure 1b. The FI undergoes large bending and extensional fluctuations. **(b)** Snapshot of the FI after 250 ns of the AA simulation. The RFH and HR2 domains (highlighted) show secondary structural plasticity. Relative to the prefusion structure, the RFH stem helices splayed into an inverted tripod that behaves as a mechanical suspension system for the BH and GP domains and the HR1-CH backbone. The HR2 secondary structure is dynamic. Bending of the FI stretched the outermost HR2 helices, triggering partial conversion into unstructured sections.

### Three hinges endow the fusion intermediate with high flexibility

Next we measured longer time FI dynamics using MARTINI CG simulations which fix the secondary structure but access timescales two orders of magnitude beyond those accessible with AA.

In 40 µs total running time over 5 runs, the FI exhibited large configurational fluctuations as in the AA simulation, bending and reorienting over a wide range of angles (Figures 3a,c). To quantify the flexibility we measured the curvature statistics along the FI (Figures 3b, S3, Supplementary Movies 2,3 and Methods). This procedure identified three high flexibility hinge regions in the base, with similar mean magnitudes of curvature and large fluctuations (Figure S4). Mapping back to the atomistic structure located the hinges as unstructured loops at the BH/RFH, RFH/HR2 and HR2/TMD interfaces, respectively (Figures 3b,c).

**Figure 3.**
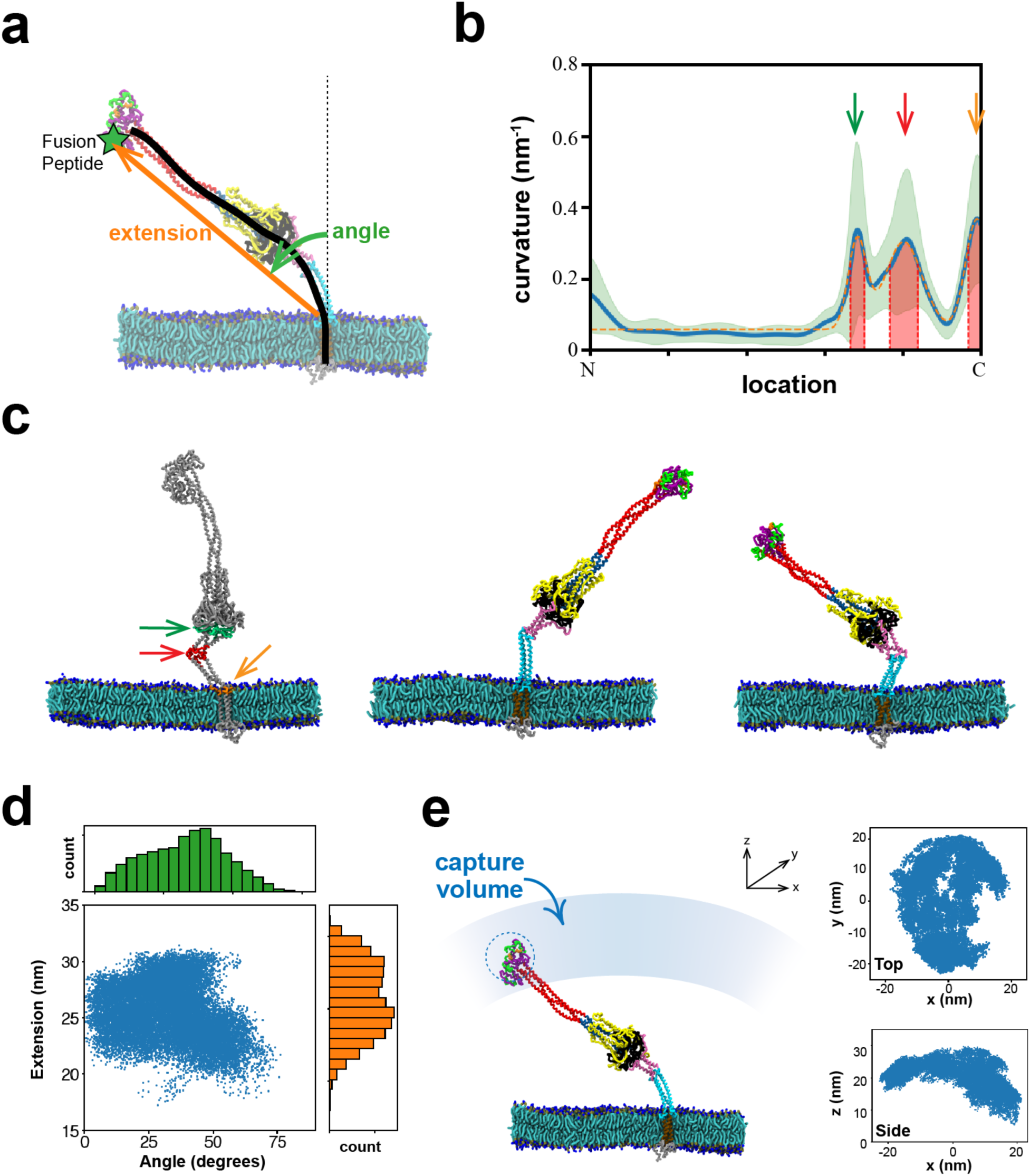
The fusion intermediate is highly flexible and visits a large capture volume. **(a)** In coarse-grained MARTINI simulations the FI had large configurational fluctuations, measured by the extension and angle of orientation of the FI backbone (black curve). **(b)** Time averaged backbone curvature versus normalized backbone arclength. Three high curvature hinges are apparent (arrows). Each hinge region (red) was defined as the quarter width of a fitted Gaussian (orange). Green envelope indicates SD. **(c)** Simulation snapshots with the three hinges highlighted, identified as residues 1084-1138, 1156-1178 and 1204-1213. **(d)** Distributions of FI extensions and angles. **(e)** The FI has a large capture volume. Top and side views of FP locations visited. The FI extension and orientation ranges are ∼21-30 nm and of ±56°, respectively (95% of sampled values) so that a large capture volume is swept out over time, shown schematically (left). Dashed circle: approximate region explored by the FP in 1 µs. **(b), (d), (e)** Statistics are averages over the last 4 µs of five 8 µs runs, for a total of 20 µs simulation time.

These hinges have roughly the same locations as three hinges identified in the prefusion CoV-2 S protein by a study combining cryo-ET and MD simulations^28^. Thus we adopt the “ankle, knee, hip” notation of that study. However, the hip hinge (RFH/BH interface) in the FI structure is more flexible than the prefusion hip due to the significantly altered RFH structure with splayed stem helices (Figure 2b).

### Large fluctuations of the fusion intermediate lead to a large membrane capture volume

The first task of the unleashed FI is thought to be capture of the host target membrane by insertion of the fusion peptides at the protomer N-termini. Due to its flexibility the FI extension ranged from ∼ 21-30 nm (mean ∼ 26 nm), its orientation varied over angles ∼ ±60° to the membrane normal, and the FPs in consequence swept out a volume ∼ 25,000 nm^1^ at rate ∼ 750 nm^1^ µs^23^ (Figures 3d,e, 95% of sampled locations).

Thus, due to the flexible base hinges combined with the large reach of the HR1-CH backbone, the FI accesses a substantial capture volume, equivalent to that of a ∼ 36-nm-diameter sphere. Following dissociation of S2 from the S1/ACE2 complex, this may help the virus rapidly reconnect with the host cell and limit refolding back into the virion membrane in a postfusion configuration without host cell contact. Indeed, postfusion spike proteins were observed by cryo-ET on intact SARS-CoV-2 virions^41^.

### Structure and dynamics of the membrane-bound fusion peptide

We used a multiscale approach to study the secondary structure and spatiotemporal statistics of the membrane-bound FP removed from its host FI (Figure 4a). The FI model of Figure 1b assumed the prefusion FP, but once bound its structure likely changes in the radically altered membrane environment. Thus we used CG MARTINI MD to bind and equilibrate the bound FP (24 µs total), followed by 2 µs of AA simulation to realistically evolve the bound secondary structure (Figure 4b).

**Figure 4.**
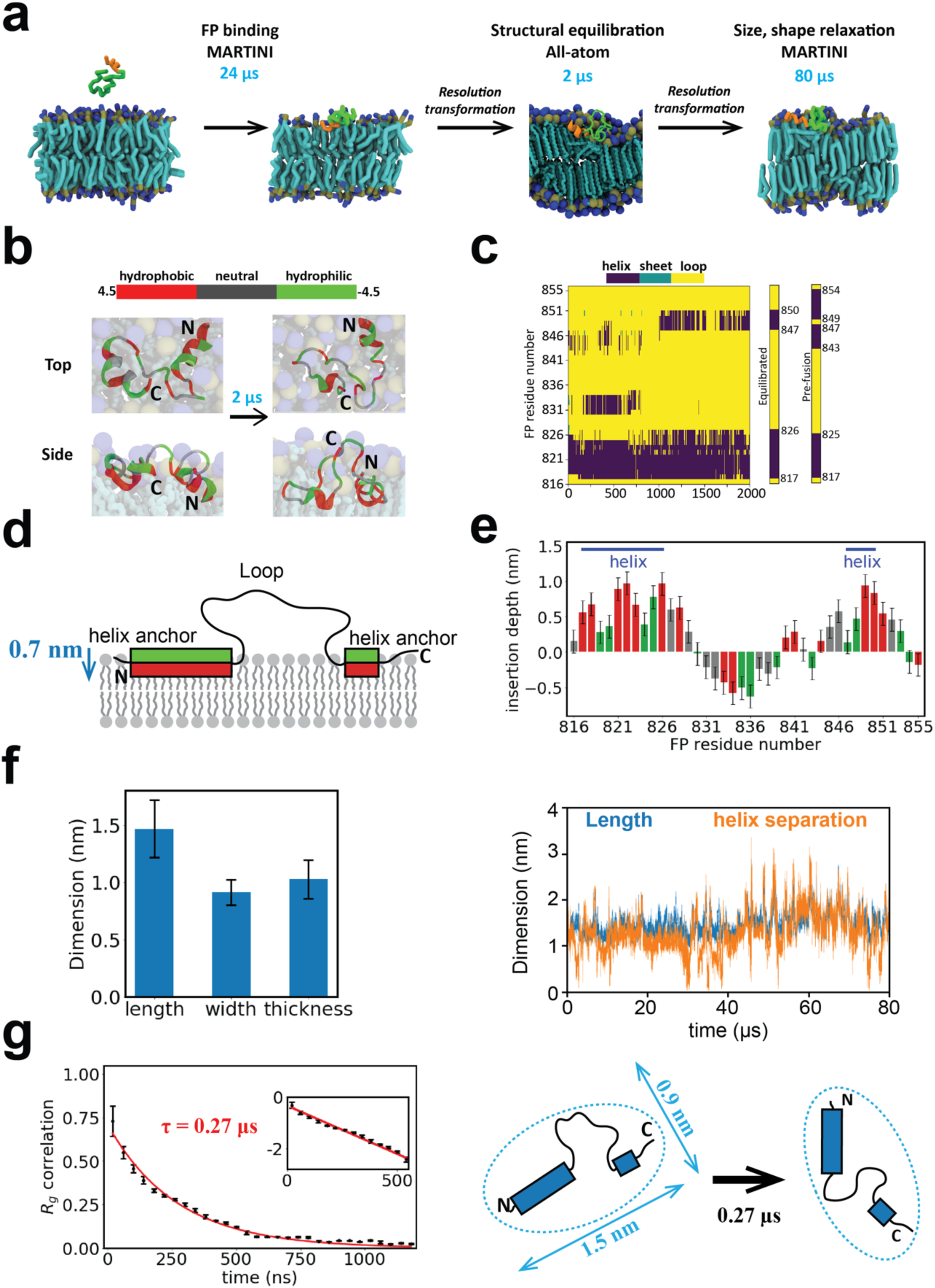
Multiscale simulations of the membrane-bound fusion peptide. **(a)** Multiscale simulation strategy to measure secondary structure and spatiotemporal statistics of the membrane-bound FP. The prefusion structure is coarse-grained to MARTINI representation and bound to and relaxed within the membrane in a 24 µs simulation (binding required ∼ 4 µs). Following backmapping to atomistic resolution, the secondary structure of the membrane-bound FP is relaxed in a 2 µs AA simulation. Assigning each residue its most frequently visited secondary structure during the final 0.8 µs of the AA simulation, the FP is again coarse-grained and its spatial dimensions and relaxation time measured in an 80 µs CG simulation. **(b)** Evolution of FP structure during the 2 µs AA simulation of (a). Initial and final states are shown. FP resides in one of three colors depending on the hydrophobicity. **(c)** Evolution of bound FP secondary structure during the 2µs AA simulation of (a). The initial (prefusion) and final (equilibrated) structures are compared. For each residue the equilibrated structure shows the most frequently adopted in the final 0.8 µs. **(d)** Equilibrated bound FP following the AA equilibration of (a), schematic. The principal anchor is the amphiphilic N-terminal helix, with a secondary amphiphilic C-terminal helix anchor. Hydrophobicity color scheme as for (b). **(e)** Mean membrane insertion depth profile along the bound FP in the equilibrated structure represented in (c) (see Methods). Mean values over 0.8 µs. **(f)** Length and helix separation of the bound FP during the 80 µs MARTINI simulation of (a). Mean dimensions averaged over the final 78 µs (left). **(g)** Temporal correlation function of the radius of gyration of the bound FP yields shape memory time *τ* = 269 ± 1 ns . (Bin size, 40 ns. 100 data points per bin.) Inset: log-lin representation. Dashed lines: exponential fit. Top view, schematic (right). All error bars: SD.

During the AA simulation the secondary structure evolved (Figures 4b, c and Supplementary Movies 4,5). The N-terminus helix barely changed, but the two C-terminal helices merged into one. (The mean total helix content was 35%, compared to ∼20% from circular dichroism spectroscopy^42^, a difference possibly explained by the differing simulated and experimental membrane compositions.) The equilibrated helices were amphiphilic and anchored the FP to the membrane with hydrophobic and hydrophilic residues oriented towards and away from the membrane, respectively (Figures 4d,e).

In an 80 µs CG simulation we then measured the statistics of the bound, equilibrated FP (Supplementary Movies 6,7). The depth of residues decreased somewhat (Figure S5a), and the C-terminal helix became repeatedly unanchored (Figure S6). The bound FP had rms length ∼ 1.5 nm and width ∼ 0.9 nm, defined as the greater and smaller of the gyration tensor eigenvalues in the x-y plane, while the rms thickness was ∼ 1.0 nm (Figures 4f, S7 and Methods). The FP extended with the anchored helices at either end, roughly speaking, as the length was strongly correlated with their separation. The radius of gyration autocorrelation function revealed a configurational memory time of ∼ 270 ns (Figure 4g), with similar times for the length, width and thickness (Figure S8).

### Measurement of fusion peptide-membrane binding rate constant

The target membrane is captured by insertion of the fusion peptide. To quantify the binding kinetics, we removed a FP from its FI host and measured the binding rate constant between two membranes separated by *h* = 5.5 nm (Figure 5). The binding time itself is not an invariant quantity, as it depends on the proximity of the membranes.

**Figure 5.**
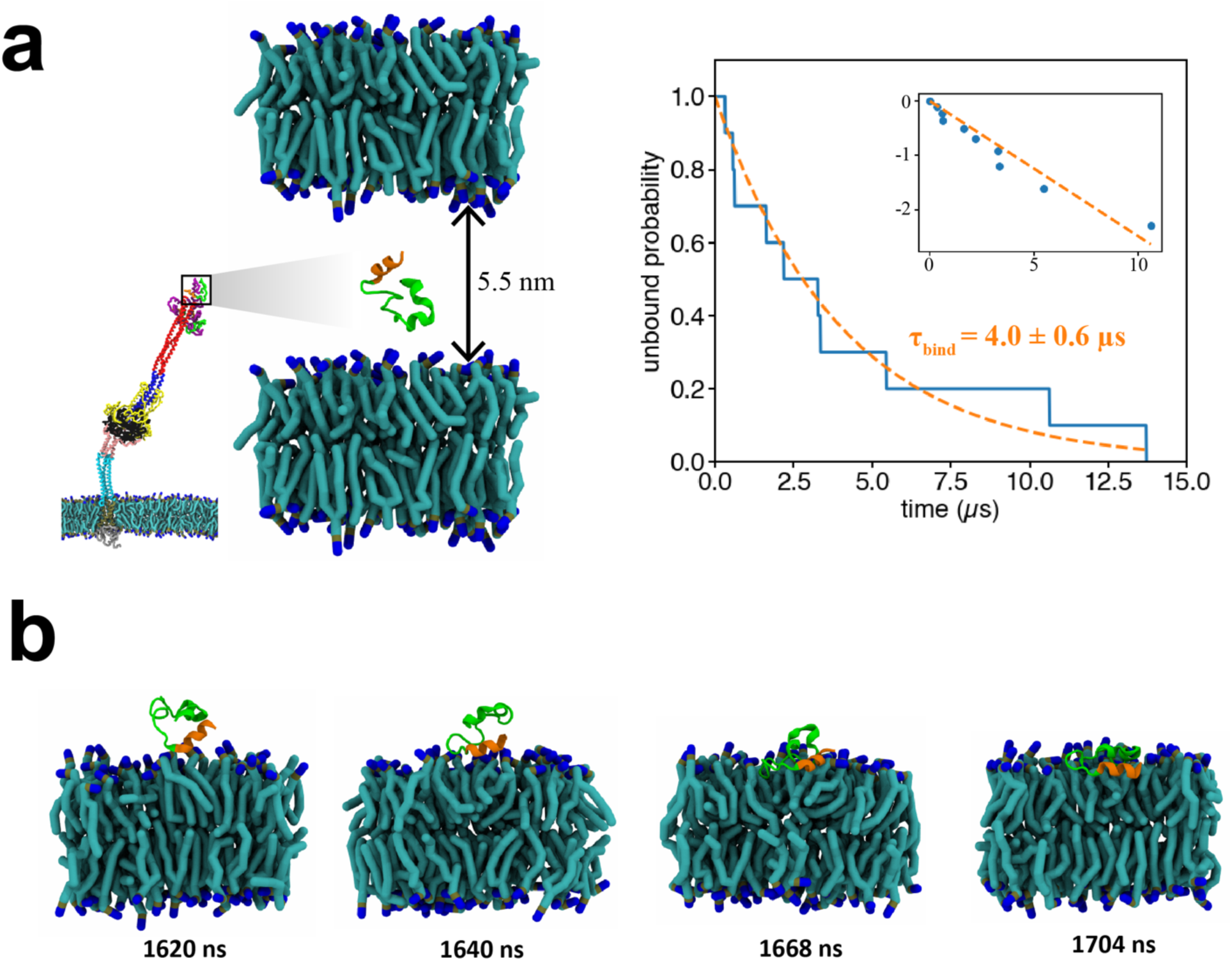
Membrane binding kinetics of an isolated fusion peptide. **(a)** Binding assay to measure the membrane binding rate constant, 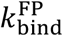, of a FP removed from its host FI. Initially the FP is positioned between two membranes separated by 5.5nm (left). FP dynamics are simulated using the CG MARTINI force field and the time to irreversibly bind the membrane is measured. The unbound fraction (blue, right) among ten simulated FPs decays exponentially with time constant *τ*_bind_ = 4.0 ± 0.6 µs (dashed orange curves). Inset: log-lin representation. **(b)** Typical binding event. The N-terminal helix (orange) is the first binding contact. To show secondary structure, the FP was back-mapped to all-atom representation.

Defining a collision as an approach to within the rms FP end-to-end distance, *R_FP_*∼ 1.6 nm, the FP collided ∼ 27 times per µs with the membrane before irreversibly binding (Figure S9). Thus, effects of initial condition dependence and diffusion-control were negligible^43^. Averaged over 10 CG MARTINI simulations the unbound probability decayed exponentially with time constant τ_bind_ ∼ 4.0 ± 0.6 µs (Figure 5a). The binding rate constant *k*_bind_ is defined by an imagined situation with a solution of FPs at density *c_FP_*contacting a membrane, such that *dρ/dt* = *k*_bind_ *c_FP_* where *ρ* is the areal number density of bound FPs^43^. From the binding assay,

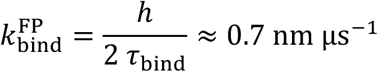

where the factor of 2 reflects the two membranes.

Importantly, binding was mediated by the N-terminal helix of the FP, which provided first contact with the membrane during a binding event (Figure 5b and Supplementary Movie 8). Since cleavage at the S2′ site would expose this helix, this is consistent with this cleavage being required for viral entry^5^.

### The fusion intermediate captures target membrane on a millisecond timescale

Next we studied membrane capture by the full FI (Figure 6a). Surprisingly, membrane binding was so much slower than suggested by the binding kinetics of the removed FP (Figure 5) as to be unobservable on available computational timescales. However, we observed binding of a truncated FI, from which we inferred a membrane capture time by the full FI of ∼ 2ms.

**Figure 6.**
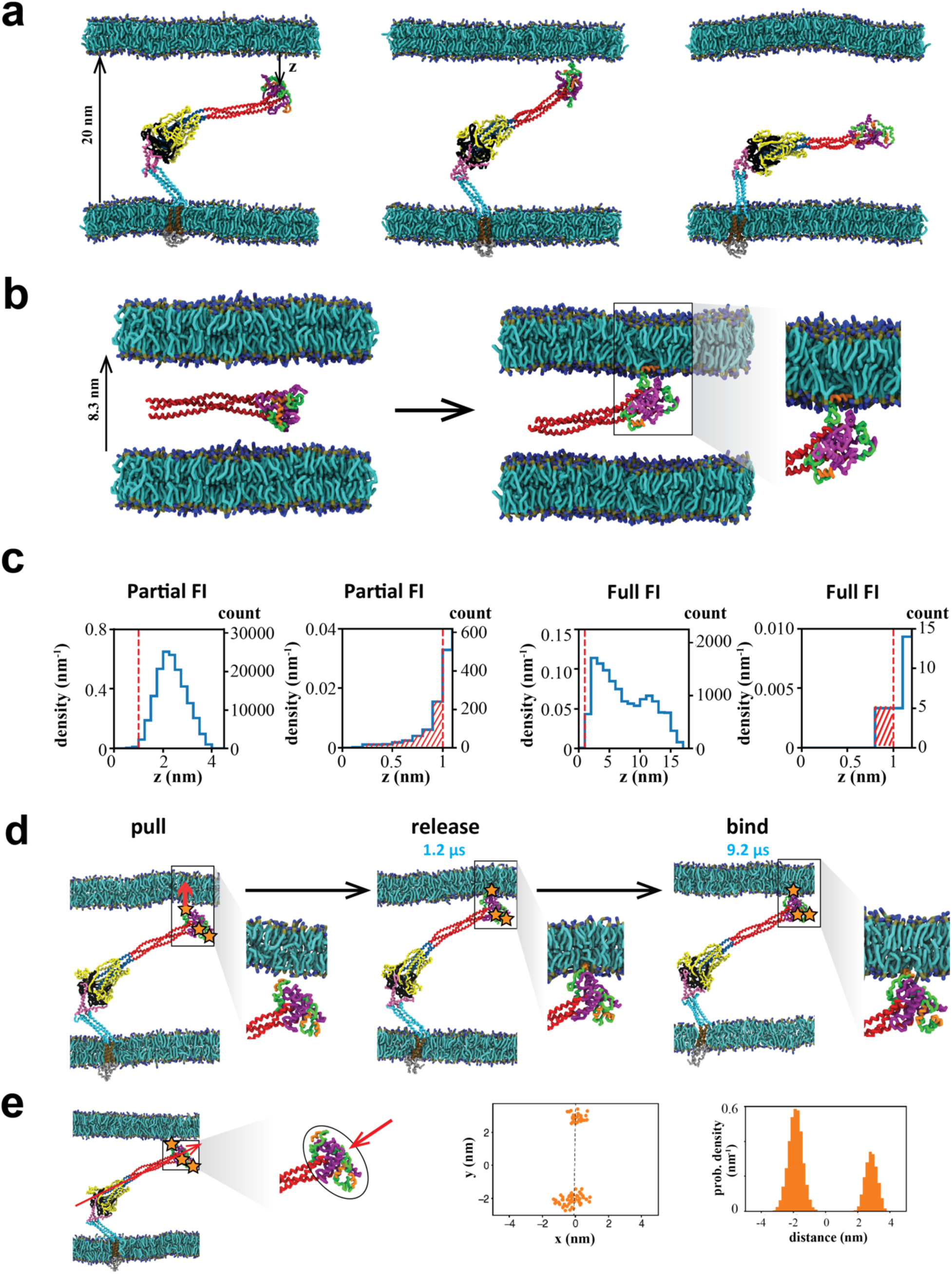
Interaction of the fusion intermediate with a target membrane. **(a)** Snapshots from CG MARTINI simulations of the full FI in the presence of a target membrane 20 nm from the viral membrane. The N-terminal helices of the FPs are shown orange. **(b)** Simulation of membrane binding by a truncated FI consisting of the HR1, CR and FP domains between two target membranes. Initial condition (left). Binding was mediated by the N-terminal FP helix (right). **(c)** Probability density versus distance *z* of the nearest N-terminal FP helix from the membrane during simulations of membrane binding by the partial or full FI. In both cases the density is depleted close to the membrane. The net probability for the N-terminal helix of the FP to lie within 1 nm of the membrane was 0.33% for the partial FI and 0.07% for the full FI (hatched areas). **(d)** Enforced binding of a FP. The FP N-terminal helix was pulled into the membrane over a period of 1.2 µs, and then released. The FP remained bound to the membrane for all of a 8 µs CG MD simulation. **(e)** The three FP-CR domains organize into a disordered laterally extended blob at the N-terminal end of the FI backbone, the FI head. Two FP N-terminal helices reside at one end of the head, one at the other end (orange stars). End view (perspective of red arrow) of N-terminal helix beads and their density distribution along the principal axis (dashed black line) in the plane normal to the backbone.

We simulated the full FI in the presence of a target membrane 20 nm from the viral membrane (Supplementary Movie 9). Enabled by its high flexibility hinges, the FI adopted highly bent shapes in which the FPs were oriented toward the membrane (Figure 6a). However, we recorded no FP-mediated binding events during a total ∼ 300 µs of CG simulation over 20 independent runs (see Methods). Thus, binding is much slower when the FPs are attached to the FI. The FP-only binding kinetics are unrepresentative, as they suggest the FI will bind at rate ∼ 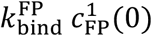 where 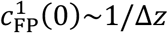 is the 1D FP density evaluated at the membrane and Δ*z*∼ 10 nm is the spread of FP N-terminal helix distances from the membrane (Figure S10). This yields a binding time 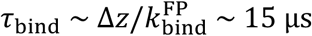, clearly a huge underestimate.

It is unclear if binding is computationally accessible even with CG dynamics, given that longer than ∼ . 3 ms is required. Thus we accelerated the kinetics by truncating the FI, excluding all domains downstream of HR1. We measured the binding time of this partial FI, consisting of HR1, CR and FP domains only, between two membranes separated by 8.3 nm (Figure 6b and Supplementary Movie 10). In two of six runs each lasting 160 µs, the membrane was captured by insertion of the N-terminal helix of one FP, after 23 µs and 133 µs. (In another case, binding was followed by unbinding after ∼42 µs.) This implies a best estimate of ∼390 ± 280 µs for the mean time for the partial FI to bind in this assay (see the Supporting Information, SI).

To translate this result to binding of the full FI, we measured the fractions of time for which one of the three FP N-terminal helices lies within 1 nm of the membrane. The full FI satisfied this criterion ∼ 5-fold less frequently than did the partial FI (∼0.07% vs. ∼0.33%, Figure 6c and Methods), suggesting membrane binding is ∼ 5-fold slower than in the partial FI assay. Thus we estimate the target membrane is captured by the full FI after ∼ 2 ms.

Finally, to verify the full FI is capable of maintaining a bound state, we enforced binding by pulling the FP of an FI into the membrane (Figure 6d). The FP remained stably bound for all of an 8 µs simulation.

### Fusion peptides and connecting regions form a disordered cluster

These results show that the ability of the FP to access target membrane is strongly constrained by its local environment in the FI. In the CG simulations this environment was a disordered cluster that the FPs and neighboring CR domains organized into, laterally extended at the N-terminal end of the HR1-CH backbone (Figure 6e). We call this the head of the FI. Two of the three FP N-terminal helices resided at one end of the head, and one more exposed helix at the other. With the host membrane ∼ 20 nm away, likely imposed by the earlier S1-ACE2 binding episode, the FI is severely bent (Figure 6e). The lateral orientation of the head appears optimal for presenting the helices to the membrane for binding in this bent configuration.

## Conclusions

The outbreak of Covid-19 saw rapid efforts to characterize the pre- and postfusion SARS-CoV-2 spike protein structures^3, 9, 11, 12^, but the structure of the fusion intermediate (FI) that facilitates fusion and entry remains unknown and the pathway to fusion and entry is poorly characterized. The first step on this pathway is capture of the host cell target membrane by the FI, but the mechanism and timescale are unknown for CoV-2 or indeed any coronavirus.

Here we built a full length model of the CoV-2 FI, extrapolating from pre- and postfusion structures^9^ (Figure 1a). From coarse-grained simulations we inferred a FI-mediated membrane capture time of a few ms. Macroscopically this is fast, suggesting that therapeutic strategies targeting the FP are limited by a small ∼ ms window during which the FP is exposed, and that targeting the refolding process^12, 21, 44^ may be more fruitful. However, a ms timescale is very long from a computational perspective: given our computational resources, membrane capture would require several hundred years of atomistic simulation (see Methods), and so is observable only with coarse-grained MD methods. These membrane binding rates were overestimated ∼ 2 orders of magnitude by simulations with the FP removed from its host fusion protein, although a 10 residue N-terminal helix directing and maintaining FP binding was identified (Figure 5b) in accord with a recent study^30^. Simulations of isolated viral fusion peptides are commonly implemented^30–32, 45^, but our results suggest they should be interpreted with caution as the fusion peptide environment is radically altered when attached to its host fusion protein.

Atomistic and coarse-grained simulations presented a picture of the FI as a machinery designed to efficiently capture membrane (Figure 7). The FI suffered large bending and tilting fluctuations due to 3 highly flexible hinge regions (Figures 2, 3, 6), similar to 3 hinges identified in the prefusion structure that were proposed to aid receptor binding^28^. We suggest the hinges are most critical to the FI. Large fluctuations may aid capture of host cell membrane by enlarging the region accessible to the FPs at the FI terminal (Figure 7c), and may help to coordinate capture by multiple FIs at different distances. Indeed, influenza, parainfluenza and HIV-1 appear to use several FIs ^14, 16, 46^. Further, by allowing the ∼25 nm long FI to bend significantly, the extreme flexibility may facilitate the prefusion-to-FI transition even in the confined circumstances of a nearby host cell membrane (Figure 6a) and allow the FI to tilt its head and present the FPs directly to the target membrane (Figures 6e and 7c). A milder flexibility was observed in the prefusion HA of influenza, which was reported to bend through ∼25^°^ mediated by a linker between the ectodomain and TMD^47^.

**Figure 7.**
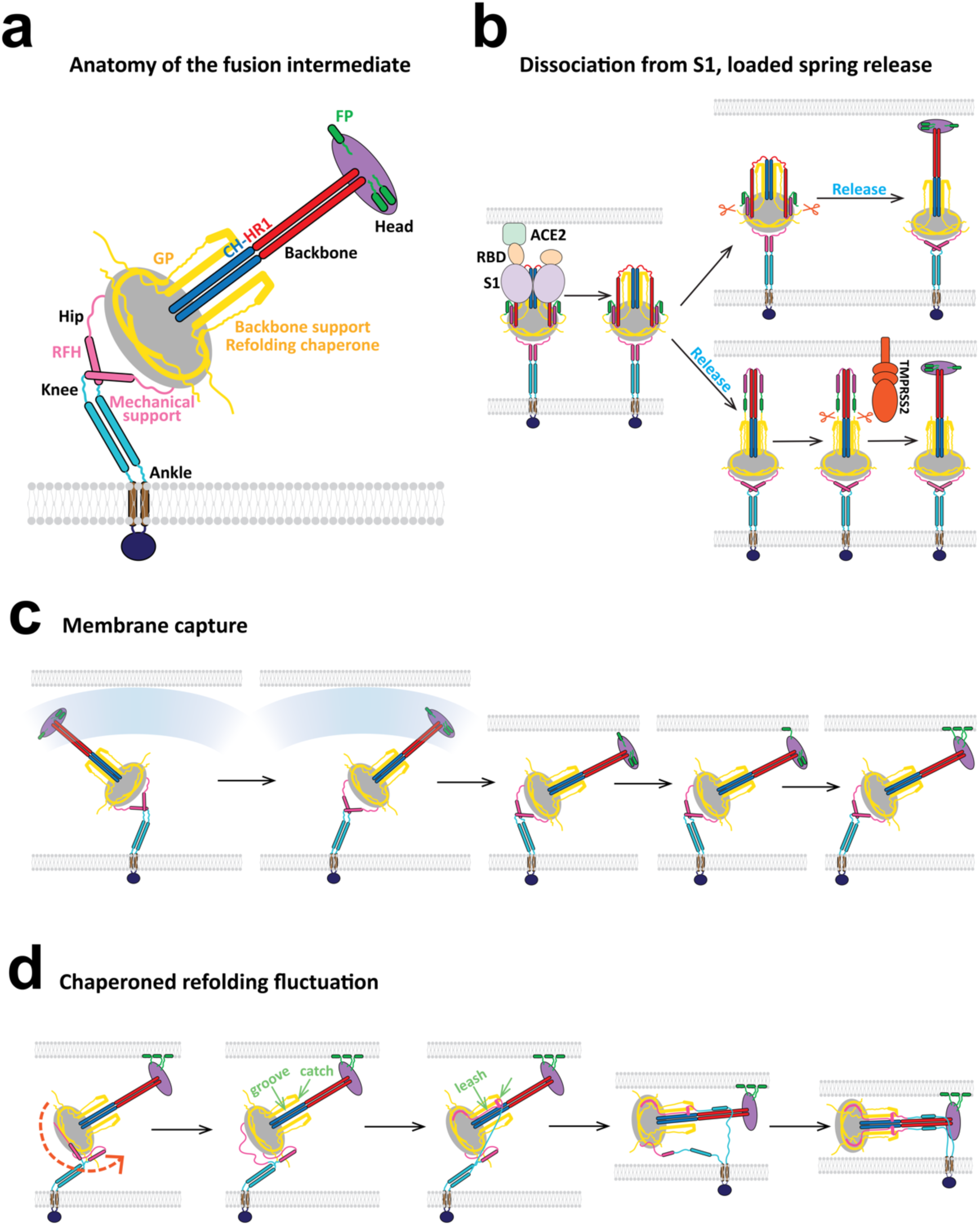
Model of the SARS-CoV-2 fusion intermediate and the pathway to fusion. **(a)** Schematic of the fusion intermediate. The ankle, knee and hip hinges impart high flexibility to the FI. RFH is an inverted tripod suspension system buffering longitudinal backbone fluctuations. GP supports the backbone and chaperones refolding. The CH-HR1 backbone provides mechanical strength and reach. The FP-CR head houses the fusion peptides for host membrane capture. **(b)** Pathway to the fusion intermediate. Following dissociation of S2 from the S1/ACE2 complex, a loaded spring release mechanism generates the fusion intermediate after proteolytic cleavage at the S2′ site (upper pathway) or before cleavage (lower pathway). RBD, receptor binding domain of S1. TMPRSS2, transmembrane protease serine 2. **(c)** Schematic of host cell membrane capture by the fusion intermediate. Three base hinges endow the fusion intermediate with high flexibility and large configurational fluctuations, so the N-terminal fusion peptides sweep out a large volume for membrane capture. **(d)** Model of fluctuation-triggered, GP-chaperoned refolding. A sufficiently large rotational fluctuation at the RFH/BH hip joint unfolds a RFH stem helix into an unstructured loop. The loop is grabbed by a GP *β*-strand in BH, initiating RFH refolding, and guided into a GP-CH groove which it packs as a leash. Leash zippering into the groove is stabilized by the GP catch, preventing unzippering. Refolding of the HR2 leash completes refolding of one protomer, pulling the membranes together and helping the other protomers refold. The trans postfusion structure catalyzes membrane fusion in cooperation with other refolded fusion proteins.

Following binding of the FI to the target membrane on a ms timescale, the next step on the pathway to fusion and cell entry is refolding of the FI that pulls the viral and target membranes together. What sets the timing of refolding? Refolding requires a major structural transition of the RFH domain, which folds into the GP domain and the CH-HR1 backbone (Figure 1a). In AA and CG simulations the RFH domains had a highly dynamic structure, permitting large bending fluctuations of the hip hinge at the RFH/BH interface (Figures 2 and 6a). This suggests the refolding time may be the waiting time for a rotational hip hinge fluctuation sufficiently large to destabilize one of the splayed RFH helices into an unstructured loop (Figure 7d). The loop would be highly susceptible to GP-chaperoned refolding. FI refolding and the drawing together of the host and viral membranes may be mutually reinforcing elements in a cooperative process, as a smaller membrane separation presumably favors refolding, while refolding decreases the membrane separation.

As the machinery that achieves cell entry, the FI is a natural therapeutic target. A number of candidate drugs have targeted the refolding step. HR2-derived peptides inhibit fusion by SARS-CoV-2 and MERS-CoV, presumably blocking formation of the HR1-HR2 six-helix fusion core^12, 21, 48^. Their efficiency as fusion inhibitors is insensitive to mutations in the spike protein^25^, suggesting potential as robust antiviral drugs.

Understanding the mechanisms of such candidate drugs and discovery of new FI-targeting drugs will be helped by establishing the structure and dynamics of this elusive intermediate. For example, another potential target is the golden peptide (GP) domain extending from the S1/S2 to the S2′ cleavage site (Figure 1). In addition to its structural role as a stabilizing socket for the CH-HR1 backbone (Figure 7a), GP chaperones RFH refolding (Figure 7d). First, GP helps initiate RFH refolding by providing the β-strand that the RFH N-terminus loop folds onto as a parallel β-strand. Second, GP supplies a groove with the neighboring CH helix into which the RFH leash packs, continuing refolding. Third, a small RFH helix is pinned by the GP catch, a U-shaped sequence including 2 small helices, that may rectify zippering of the RFH leash by preventing its unravelling from the groove. Thus, GP- or RFH-derived peptides could inhibit FI refolding by binding the RFH or GP domains. Such peptides might also stabilize the short-lived unfolded FI for visualization.

Interestingly, a recent study identified an 8-residue region in the prefusion RFH stem helix as the epitope of two cross-reactive monoclonal antibodies^49^. This region becomes the small RFH helix that engages the GP catch during refolding, together with the four downstream residues. Thus, the antibodies may neutralize CoV-2 by binding RFH and interfering with the GP catch that rectifies RFH refolding. Our simulations suggest another possibility is that binding of the RFH stem helices in the FI alters their dynamics and lowers the hip hinge flexibility, with possible consequences for membrane capture and/or refolding. This would be similar to the effect of antibodies targeting the linker domain adjacent to the TMD of HA, which reduced the linker flexibility and suppressed orientational fluctuations ^47^.

In summary, the extended FI is the fusogenic form of the spike protein that captures host cell membrane for fusion and entry and is a critical but relatively unexplored therapeutic target. Our model suggests a loaded-spring mechanism generates the FI from the prefusion structure, related to the mechanism for HA of influenza^15, 37^ . The FI has unexpectedly large bending fluctuations that help it capture membrane in a few ms and may trigger the refolding transition that draws the viral envelope and host membranes together for subsequent fusion (Figure 7). These results provide an account of a critical episode during cell entry and offer a framework for rational design of new therapeutic strategies to disable the FI.

## Methods

### Building a complete structure for the fusion intermediate of the SARS-CoV-2 spike protein

The primary sequence of the SARS-CoV-2 S protein was obtained from the NCBI database (GenBank: MN908947). We first built the HR1-RFH portion of the FI (including the associated GP domain). We used Modeller^50^ with two specified templates: the postfusion structure (PDB: 6XRA) for HR1-BH and GP, and the prefusion structure (PDB: 6XR8) for RFH. Several constraints including 3-fold symmetry and preservation of secondary structure were specified. The C-terminal domains (HR2, TMD and CT) were then appended to the HR1-RFH portion one by one using Modeller. The source for HR2 was a SARS-CoV HR2 NMR structure (PDB: 2FXP), while the TMD/CT structure was predicted by the QUARK server by providing the primary sequences. The FP and CR were then extracted as a whole from the prefusion structure (PDB: 6XR8) and appended to the HR1 domain in the FI with an arbitrary angle using Pymol. Finally, a complete GP structure was made in the FI using Modeller, by appending the N-terminal portion (residues 686-702) and C-terminal portion (residues 771-815) of the prefusion structure to the solved postfusion GP (residues 703-770).

### Multiscale molecular dynamics simulations

To simulate the FI and its FP we used two complementary molecular dynamics (MD) approaches, all-atom (AA) and coarse-grained (CG) MARTINI. Pure DPPC membranes represented the viral envelope and the host cell membrane for simplicity. All simulation systems were solvated with 150 mM NaCl at neutral pH. Each simulation system was energy-minimized and equilibrated before the production run, which was performed in the NPT ensemble at 1 bar and 310 K using GROMACS 2019.6^51, 52^. The AA simulation of the FI (Figure 2) required one day of computation to run the FI for 10 ns. At this rate, ∼ 500 years would be needed to achieve the ∼ 2 ms timescale of membrane capture (Figure 6c). For details of the simulations and the analysis see SI.

### All-atom simulations

For the FI simulation (Figure 2), the FI model structure (Figure 1b) was placed in a planar lipid bilayer in a 16 × 16 × 43 nm^1^ box using the CHARMM-GUI membrane builder^53^. The production simulation was run for 406 ns using the CHARMM36 force field^54, 55^. Secondary structure was identified using the *dssp* algorithm^56, 57^.

To simulate the membrane-bound FP (Figure 4b), the final configuration of the FP and the membrane to which it was bound in a given CG FP binding simulation was converted to atomic resolution in CHARMM36 force field^54, 55^ using the *backward* utility^58^. The production simulation was run for 2 µs. The membrane insertion depth of each FP residue was defined as the vertical distance between the residue center of gravity (COG) and the COG of the PO_V_ groups in the membrane leaflet to which the FP was bound.

### Coarse-grained simulations

All running times reported here are 4 times the raw MARTINI running time, to compensate for the faster sampling of the MARTINI model^59^. Time-dependent data in MARTINI simulations were analyzed after adjusting in this fashion.

For the FI simulations in the absence of a target membrane (Figure 3), the FI model structure (Figure 1b) was first mapped into the MARTINI CG representation using the *martinize* utility^60, 61^ and then placed in a planar membrane in a 30 × 30 × 50 nm^1^ box using the *insane* utility^62^. Five production simulations starting from the same equilibrated system were run for 8 µs.

For the isolated FP-membrane binding assay (Figure 5), the atomistic structure of the FP was extracted from the prefusion cryo-EM structure (PDB: 6XR8) and coarse-grained into MARTINI representation. Then the FP was placed ∼1 nm above a planar bilayer in a 7 × 7 × 10 nm^1^ box. By implementing periodic boundary condition, this is equivalent to using two planar membranes separated by ∼5.5 nm. The C-terminal carboxyl group was neutralized as it connects to the CR in the full-length FI. Ten 24 µs parallel runs were performed. Binding events were identified by comparing the COGs of the FP and the PO4 beads in each membrane leaflet (see SI).

To simulate an equilibrated FP bound to a membrane (Figure 4), the final configuration of the FP and the membrane to which it was bound in the AA simulation was coarse-grained into MARTINI representation The production simulation was run for 80 µs. In a variation of this procedure, the FP C-terminal helix was transformed to an unstructured loop by changing the secondary structure file of the FP, an input to the coarse-graining utility. We then measured the membrane insertion depth of each residue, gyration tensor, radius of gyration, length, and width of the FP (see SI).

For ten of the simulations of the FI interacting with a target membrane, ten snapshots from three FI simulations without membranes were used as initial conditions, in which the FI ectodomain protruded less than 20 nm normal to the membrane. Another pre-equilibrated planar membrane (run for 4 µs) was placed 20 nm above the membrane anchoring the FI and this configuration was run for 8 µs. These runs were used to calculate the probability distributions of Figure 6c. Ten additional simulations (each lasting 22.4 µs) used a biased initial condition with the COG of the nearest N-terminal FP helix within 1 nm of the membrane and the unpaired helix facing the membrane. Here, the membrane position was defined as the mean location of all PO_4_ beads in the lower leaflet of the upper membrane.

To pull a FI-attached FP into a target membrane (Figure 6d), an initial condition was chosen with the nearest unpaired N-terminal FP helix lying within 1 nm of the membrane. The helix was then pulled towards the target membrane at speed 2.5 nm/µs for ∼1.2 µs, until the COGs of the helix and membrane were separated by ∼0.1 nm in the *z* direction. The pulling force was then released, and the simulation was run for 8 µs.

For simulations of membrane binding by the partial FI (Figure 6b), the structure of the HR1, CR and FP domains in the MARTINI CG representation was extracted from the final configuration of a FI simulation. This partial FI was positioned above a planer bilayer in a 20 × 20 × 13 nm^1^ box with periodic boundary conditions, equivalent to two planar membranes separated by ∼8.3 nm. The C-terminal carboxyl group was neutralized. Six 160 µs parallel runs were performed. Binding events were identified similarly to the isolated FP assay (see SI). To infer the binding time of the full FI from the measured partial FI binding time, we assumed the binding probability per unit time for a given FP-CR-HR1 configuration is independent of the remainder of the FI.

## Data availability

All simulation data supporting the findings in this paper are available from the corresponding author upon reasonable request. The structure of the model of the SARS-CoV-2 spike fusion intermediate presented here is available at https://github.com/RuiSu11/cov-2_public

## Acknowledgements

We gratefully acknowledge high-performance computing resources from Microsoft Azure, provided through the COVID-19 HPC Consortium from government, industry and academia that volunteers computing time and resources to support COVID-19 research. Particular thanks are due to Geralyn Miller, Jer-Ming Chia and John Sawyer at the Microsoft AI for Good Research Lab for building and maintaining the HPC system. Columbia University’s Shared Research Computing Facility is gratefully acknowledged for the initial stages of this project.

## Author contributions

B.O’S designed the research and performed mathematical analysis and analysis of the structure. R.S and J.Z built the structure and performed the simulations. R.S, J.Z and B.O’S analyzed the data. B.O’S and R.S wrote the paper with contributions from J. Z.

## Supplementary Methods

This section provides more detailed information for each simulation described in Methods.

### All-atom simulation of the fusion intermediate

The full-length fusion intermediate (FI, Fig. 1b) with its TMD inserted in the membrane was placed in a simulation box of 16 × 16 × 43 nm^1^. The membrane-protein system was built using the CHARMM-GUI membrane builder^1^, consisting of 786 lipids for the pure DPPC membrane. The termini and ionizable residues were treated in their charged states assuming neutral pH. The disulfide bond in FP was added according to the prefusion structure (PDB: 6XR8), while the disulfide bonds in the other domains, BH and GP, were added based on the postfusion structure (PDB: 6XRA, the disulfide bonds were conserved in the pre- and postfusion structures). The resulting simulation box contained approximately 300,000 water molecules and was neutralized with 150 mM NaCl ions. The TIP3P model was used for water^2^.

The system was first energy-minimized for 1,000 steps. Then, 2 equilibrations in the NVT ensemble were each performed for 0.1 ns with position restraints on all protein atoms. Subsequently, 4 equilibrations in the NPT ensembles were each performed for 0.5 ns with position restraints on protein heavy atoms. The production simulation was run for 406 ns in the NPT ensemble at 1 bar and 310 K and with a time step of 2 fs. The temperature and pressure were maintained using the Nosé-Hoover thermostat^3, 4^ and Parinello-Rahman barostat^5^, respectively. All the energy-minimization, equilibration, and production simulations were performed using GROMACS 2019.6^6, 7^. The secondary structure of each residue in the FI was analyzed using the *dssp* algorithm^8, 9^ for every 0.1 ns.

#### Coarse-grained simulation of a fusion intermediate with uncleaved S2’ sites

Atomistic coordinates of the full-length fusion intermediate (Fig. 1b) were converted onto the MARTINI 2.2 topology using the *martinize* utility and placed in a simulation box of 40 × 40 × 50 nm^1^. The unsolved C-terminal part of the GP domain in the postfusion structure^10^ (residues 771-815) was forced to be a loop, by changing the input secondary structural file to the *martinize* utility. The terminal and ionizable residues were treated in their charged states assuming neutral pH. The box was then solvated by approximately 600,000 coarse-grained water particles and was neutralized by 150 mM NaCl ions.

In each protomer, the FP N-terminus residue (residue 816) and the GP C-terminus residue (residue 815) were pulled together at a constant rate of 10 nm/µs by a harmonic potential with a force constant of 500 kJ mol^23^ nm^2$^. Position constrains by a harmonic potential with force constant of 1,000 kJ mol^23^ nm^2$^ were applied to the beads in the domains other than GP, FP and CR. The pulling simulation took ∼1.5 µs so that in the final configuration, the COG distance between residues 815 and 816 in one of the three protomers reaches ∼0.5 nm.

The final coordinates of the coarse-grained FI were then backmapped into atomistic resolution in CHARMM36 force field^11, 12^. The residues 815 and 816 were covalently connected in the protomer with the smallest separation between the two residues, and this protomer was duplicated twice to make a trimer using Pymol. Now a structure of the FI with its S2’ sites uncleaved was created. Then, the all-atom structure of the FI with its S2’ sites uncleaved was converted into MARTINI coarse-grained representation using the *martinize* utility. A simulation box of 30 × 30 × 50 nm^1^ was generated using the *insane* utility, in which the coarse-grained FI with its S2’ sites uncleaved was inserted in a crystalline DPPC bilayer consisting of 2,831 coarse-grained lipids that represented the viral envelope.

The system containing the coarse-grained FI with its S2’ sites uncleaved on the viral envelope was first energy-minimized for 2,000 steps in the vacuum. Subsequently, the box was solvated with approximately 300,000 coarse-grained water particles and neutralized with 150 mM NaCl ions. The system was then energy-minimized for 2,000 steps and equilibrated for 4 ns in NPT ensemble sampling. Then the system was subjected to a production simulation lasting 4 µs in the NPT ensemble. The system temperature, membrane tension, and system pressure were maintained at the same value using the same thermostat and barostat as mentioned above.

#### Coarse-grained simulations of the fusion intermediate

Atomistic coordinates of the modeled full length fusion intermediate (Fig. 2a) were converted onto the secondary structure based coarse-grained MARTINI 2.2 topology^13, 14^ using the *martinize* utility. The termini and ionizable residues were treated in their charged states assuming neutral pH. Disulfide bonds were added to the same residues as in the all-atom simulation. The simulation box of 30 × 30 × 50 nm^1^ was generated using the *insane* utility^15^, in which the coarse-grained (CG) FI with its TMD inserted in a crystalline DPPC bilayer consisting of 3,024 coarse-grained lipids that represents the viral envelope.

The system was first energy-minimized for 2,000 steps in vacuum. Subsequently, the box was solvated with approximately 300,000 coarse-grained water particles and neutralized with 150 mM NaCl ions. The system was then energy-minimized for 2,000 steps and equilibrated for 4 ns in NPT ensemble sampling. Then the system was subjected to 5 independent production simulations, each lasting 8 µs in the NPT ensemble. Equations of motion were integrated using the Verlet leapfrog algorithm with a 80 fs time step. Bonds were constrained with the LINCS algorithm. The system temperature (310K) was maintained by the velocity rescale thermostat^16^. The membrane tension (0.05 pN/nm) and the system pressure (1 bar) were maintained by the Berendsen barostat with surface-tension coupling^17^.

### Fitting a curve to the fusion intermediate ectodomain

To fit a curve to represent the FI ectodomain backbone (residues 912-1237) in the MARTINI simulations of the FI (Figs. 3 and S2), we first represented the backbone by points. Each point represented a residue and its position was calculated by the averaged position of the MARTINI backbone bead (the non-sidechain bead) of each residue among the three protomers. The coordinates of the beads were extracted from the simulation trajectory using the *mdtraj*^18^ python package. Then the FI backbone was divided into an upper part (residues 912-1191) and a lower part (1152-1237) with an overlapping region of 40 residues. Each part was separately smoothed by the following steps. (1) All points (*x*_1_, *y*_1_, *z*_1_) were calibrated so that the center of gravity (COG) was at the origin, (2) All points were rotated so that the new x,y,z axis are aligned with the eigenvectors of the gyration tensor of the rotated points 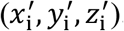. This rotation maximized the root mean square projected length onto the *z* axis. (3) A smoothed curve 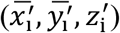 was generated by the LOWESS algorithm in python, in which the smoothed 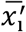 and 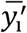 value for each 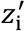 was set by its neighboring points spanning one tenth of the entire z range. Then the smoothed upper/lower part was rotated back to the original orientation and the overlapping region was averaged between the upper/lower parts. Finally the reconnected points were smoothed by the B-spline method with 4^th^ order polynomial functions. To find the location (from the N- to the C-terminus) of each domain on the fitted the curve, the curve was first reparametrized by its normalized arclength. The location of each residue was determined as the normalized arclength of the nearest point on the fitted curve to the position of this residue averaged over the three protomers, using only the backbone bead to locate the residues.

### The isolated fusion peptide binding assay

The atomistic structure of the fusion peptide (FP, residue 816-855) was extracted from the crystal structure of the prefusion structure (PDB: 6XR8). Then the atomistic structure of the FP was coarse-grained into the MARTINI representation. The ionizable residues were treated in their charged states assuming neutral pH. The C-terminal carboxyl group at residue 855 was neutralized by changing the type of the backbone bead from Qa to Na and changing the backbone bead charge from -1 to 0 in the itp topology file, as the residue 855 connects CR in the full-length FI. The disulfide bond was added according to the solved prefusion structure (PDB: 6XR8). The coarse-grained FP was placed approximately 1 nm above a crystalline DPPC bilayer consisting of 162 coarse-grained lipids in a 7 × 7 × 10 nm^1^ box using the *insane* utility^15^. By implementing periodic boundary condition, this is equivalent to placing a FP between two planar membranes separated by ∼5.5 nm.The system was first energy-minimized for 500 steps in vacuum. Subsequently, the box was solvated with approximately 2,000 coarse-grained water particles and neutralized with 150 mM NaCl ions. The system was then energy-minimized for 500 steps and equilibrated for 4 ns in the NPT ensemble with a 80 fs time step. Then the system was subjected to 10 independent production simulations, each lasting 24 µs in the NPT ensemble. The system temperature (310K) was maintained by the velocity rescale thermostat^16^. The membrane tension (0.05 pN/nm) and the system pressure (1 bar) were maintained by the Berendsen barostat with surface-tension coupling^17^. A binding event was defined to be when the z coordinate of the FP COG first had a value that positioned it below the upper membrane leaflet and above the lower membrane leaflet, where the leaflet locations were defined as the average locations of the PO4 beads in each leaflet.

#### All-atom simulation of a membrane-bound fusion peptide

In one of the CG simulations in the FP-only binding assay, the final configuration of the FP and the membrane to which it was bound is converted to atomic resolution in CHARMM36 force field^11, 12^ using the *backward* utility^19^. The ionizable residues are treated in their charged states assuming neutral pH. The C-terminal carboxyl group is neutralized. The Disulfide bond is added to the same residues as in the FP-only binding assay. The resulting simulation box contained approximately 9,000 water molecules and was neutralized with 150 mM NaCl ions. The TIP3P model was used for water^2^.

4 equilibrations in the NVT and NPT ensembles were each performed for 50 ns with position restraints on peptide heavy atoms. The production simulation was run for 2 µs in NPT ensembles at 1 bar and 310 K and with a time step of 2 fs. The temperature and pressure were maintained using the Nosé-Hoover thermostat^3,4^ and Parinello-Rahman barostat with isotropic coupling^5^, respectively. The secondary structure of each residue in the FP was analyzed using the *dssp* algorithm^8, 9^ for every 0.1 ns. The membrane insertion depth of each FP residue was defined as the vertical distance between the residue COG and the COG of PO_4 groups in the membrane leaflet to which the FP was bound.

#### Coarse-grained simulations of an equilibrated fusion peptide bound to a membrane

In the AA simulation of a membrane-bounded FP simulation, the final configuration of the FP and the membrane to which it was bound was converted to MARTINI CG representation using the *martinize* utility. The ionizable residues were treated in their charged state. The C-terminal carboxyl group was neutralized.

The system was first energy-minimized for 1,000 steps in the vacuum. Subsequently, the box was solvated with water particles and ions to attain a salt concentration of 0.15 M. The system was then energy-minimized for 5,000 steps and equilibrated for 0.8 ns in the NPT ensemble. Then the system was subjected to the production simulation for 80 µs in the NPT ensemble. The system temperature (310K) and pressure (1 bar) were maintained by the velocity rescale thermostat and Parinello-Rahman barostat, respectively.

The membrane insertion depth of each residue was defined in the same way as for the AA simulation (see above). The gyration tensor *M* of the FP was defined as

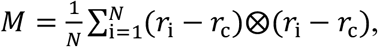

where ⊗ represents the dyadic product, *r_i_* is the coordinate of the *i*th bead in the FP, *N* is the total number of beads, and r_i_ is the COG of the FP, as 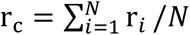. The radius of gyration *R_g_* was computed as the square root of the trace of *M*. The length and the width of the FP were defined as, respectively, greater and smaller of the eigenvalues of *M* projected onto the x-y plane, and the thickness was defined as 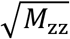.

#### Coarse-grained simulations of the fusion intermediate interacting with a target membrane

For ten of the simulations of the FI interacting with a target membrane, ten snapshots from three FI simulations (one membrane) were chosen as the initial condition in which the FI ectodomain protruded ∼20 nm normal to the membrane. Another pre-equilibrated planar membrane (run for 4 µs) was placed 20 nm above the membrane anchoring the FI in the selected configurations. Each configuration containing an FI and two membranes was re-solvated by approximately 200,000 coarse-grained water particles and neutralized by 150 mM NaCl ions. Then, each of the 10 systems was equilibrated for 4 ns, and subject to a production simulation for 8 µs in the NPT ensemble. The system temperature (310K) was maintained by the velocity rescale thermostat. The membrane tension (0.05 pN/nm) of the two membranes and the system pressure (1 bar) was maintained by the Berendsen barostat with surface-tension coupling.

In an additional set of simulations, all ten runs started from a biased initial condition, in which the COG of the nearest N-terminal FP helix was within 1 nm of the adjacent membrane and the FI head was in the (1,2) configuration. Here, the membrane position was defined to be the mean location of all the PO4 beads in the lower leaflet of the upper membrane. Each production simulation lasted 22.4 µs with the same system temperature, pressure and the membrane tension.

### Coarse-grained simulations of membrane binding by the partial fusion intermediate

The structure of the HR1, CR and FP domains in the MARTINI CG representation was extracted from the final configuration of a simulation of the FI interacting with a target membrane. The ionizable residues were treated in their charged states assuming neutral pH. The C-terminal carboxyl group at residue 984 was neutralized by changing the type of the backbone bead from Qa to Na and changing the backbone bead charge from -1 to 0 in the itp topology file, as the residue 984 connects CH in the full-length FI. This partial FI was positioned above ∼ 0.5 nm above a crystalline DPPC bilayer consisting of 676 coarse-grained lipids in a 20 × 20 × 13 *nm*^1^ box with periodic boundary conditions, equivalent to two planar membranes separated by ∼8.3 nm.

The system was first energy-minimized for 500 steps in vacuum. Subsequently, the box was solvated with approximately 26,000 coarse-grained water particles and neutralized with 150 mM NaCl ions. The system was then energy-minimized for 2,000 steps and equilibrated for 4 ns in the NPT ensemble with a 80 fs time step. Then the system was subjected to 6 independent production simulations, each lasting 160 µs in the NPT ensemble. The system temperature (310K) was maintained by the velocity rescale thermostat^16^. The membrane tension (0.05 pN/nm) and the system pressure (1 bar) were maintained by the Berendsen barostat with surface-tension coupling^17^.

The exact time for a binding event was defined to be when the vertical distance between the FP N-terminal helix COG and the membrane first had a value smaller than the averaged FP-membrane distance when the partial FI was stably bound. The membrane locations were defined as the average locations of the PO4 beads in leaflet to which the partial FI was bound.

To infer the averaged binding time *τ* of a partial FI, we assumed that at time *t* the unbound probability *P* follows an exponential function *P* = exp (−*t*/*τ*). Given that at the end of the six simulations *t* = 160 μ*s* the unbound probability is 4/6, we estimated *τ* = 390 ± 280 µ*s*, where the uncertainty of the estimation is obtained using from error propagation.

### Pulling the fusion peptide into the target membrane

An initial condition was chosen with the nearest N-terminal FP helix lying within 1 nm of the membrane and with the FI head in the (1,2) configuration. The FP N-terminal helix was pulled vertically towards the upper membrane at a constant rate of 2.5 nm/µs by a harmonic potential with a force constant of 1,000 kJ mol^23^ nm^2$^. The system temperature, membrane tension, and system pressure were maintained at the same value using the same thermostat and barostat as mentioned above. The pulling process took ∼1.2 µs. In the final configuration the distance between the COG of the FP N-terminal helix and the COG of the entire upper membrane reached ∼0.1 nm, and the FP N-terminal helix was pulled into the adjacent membrane patch by ∼1 nm. Then the force was released and a subsequent 8 µs CG simulation was run, in which the system temperature, membrane tension, and system pressure were maintained at the same value using the same thermostat and barostat.

## Supplementary Figures

**Supplementary Figure 1.**
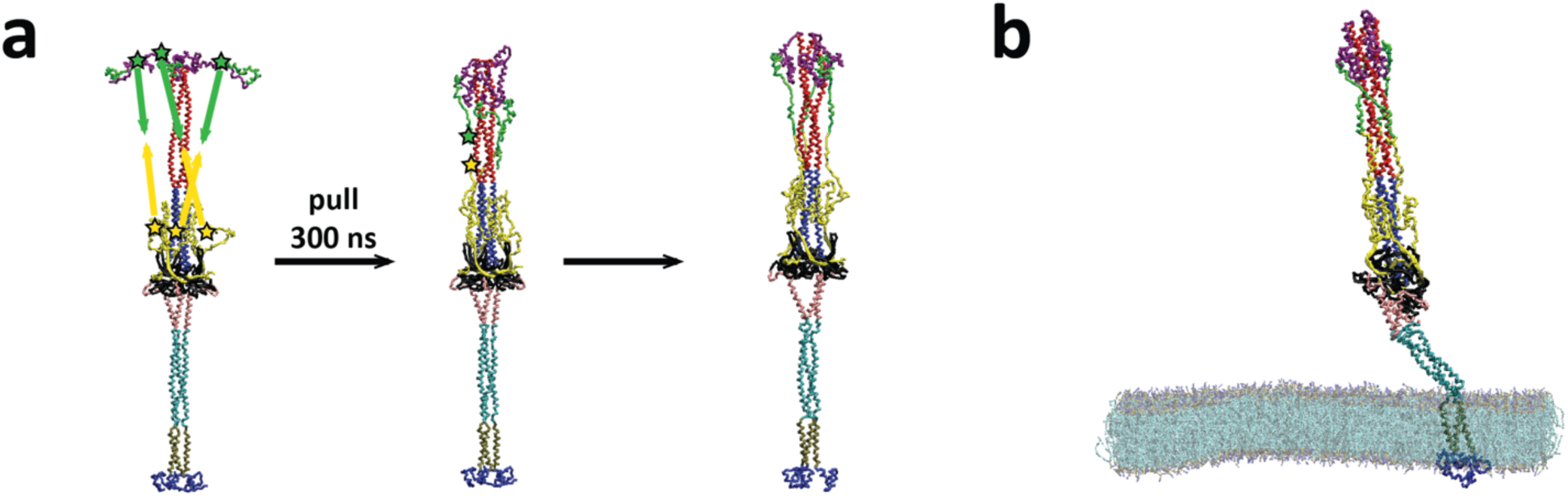
Simulation of the fusion intermediate with uncleaved S2’ sites. **(a) Construction p**rocedure for a fusion intermediate with uncleaved S2’ sites. Starting from the model structure in Fig. 2b, the C-terminus of three GPs and the N-terminus of three FPs were pulled together in ∼1.5 µs in a MARTINI CG simulation. Then the C-terminus of GP and the N-terminus of FP in one protomer were connected covalently (stars). This protomer was duplicated twice to generate a FI homotrimer with uncleaved S2’ sites. **(b)** Snapshot from a 4 µs simulation. The fusion peptides (green) are sequestered. The FI with uncleaved S2’ sites exhibits similar flexibility to the normally cleaved FI.

**Supplementary Figure 2.**
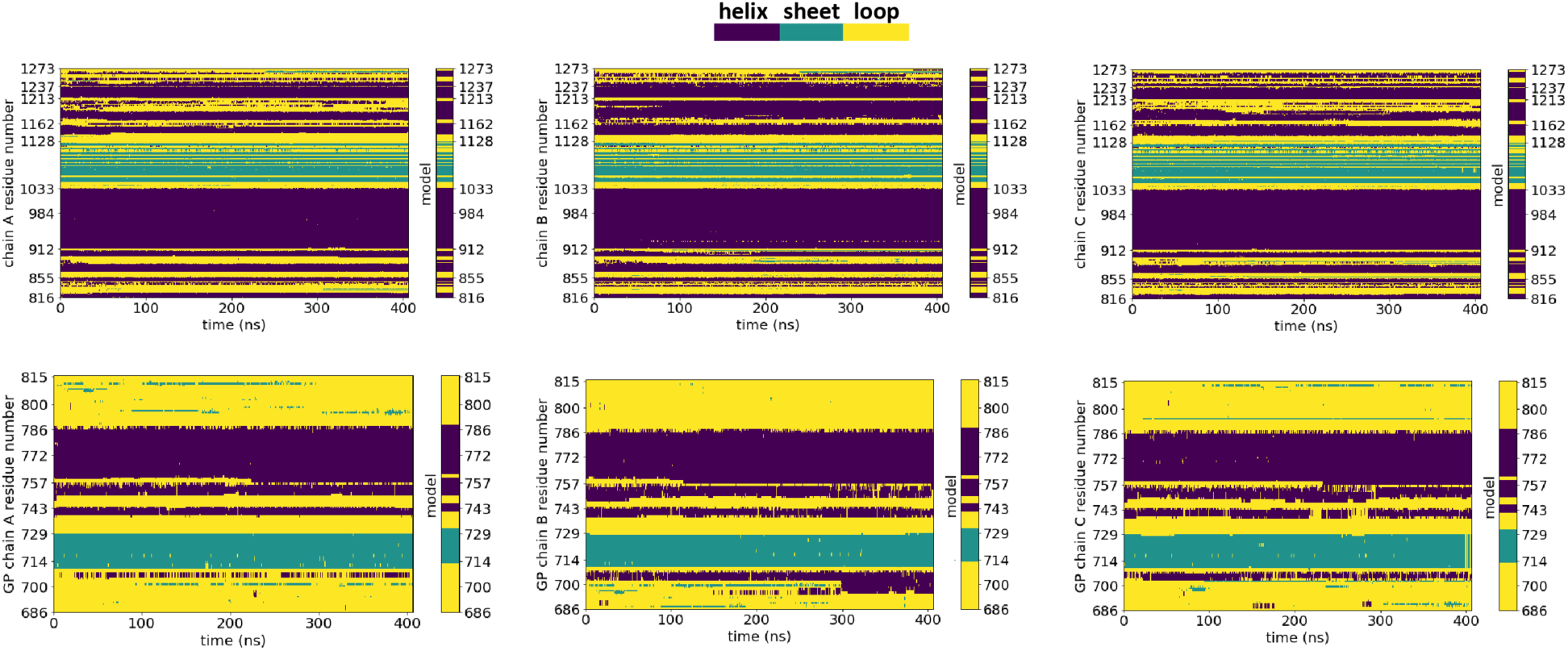
Evolution of the secondary structure of the fusion intermediate during 406 ns of all-atom simulation. The secondary structure of each residue was measured every 0.1 ns during the AA simulation (Fig. 2). Each panel refers to one protomer of either the main body of the S2 subunit (top row) or the cleaved GP (bottom row). The secondary structure of the fusion intermediate model of Figure 1b is shown to the right of each panel for comparison.

**Supplementary Figure 3.**
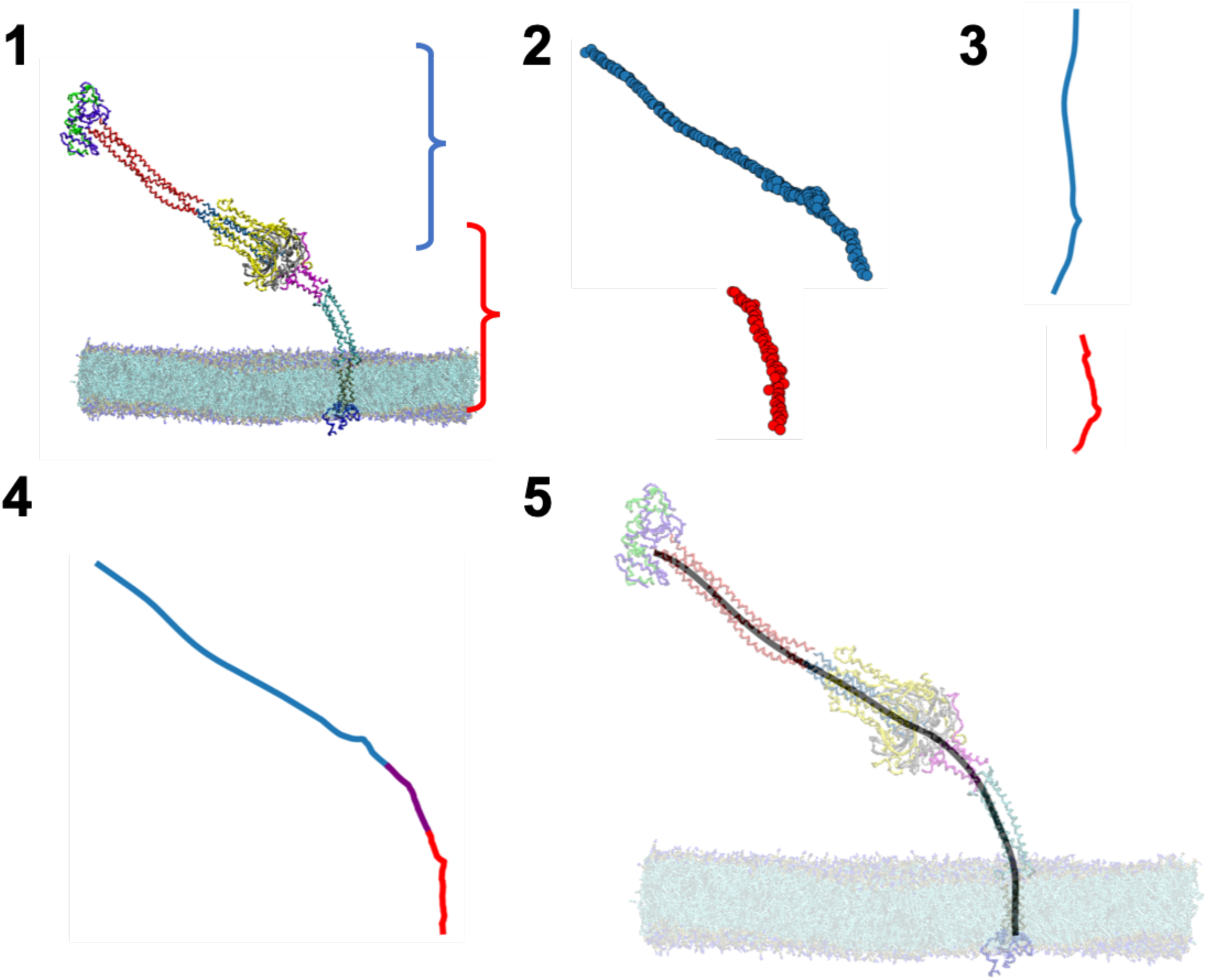
Fitting a curve to represent the fusion intermediate ectodomain. (1) The coordinate of each coarse-grained bead is extracted by Python package *mdtraj*. (2) Each residue is represented by one point, averaged over three protomers. The upper part of the FI (blue, residues 912-1191) and the lower part (red, residues 1152-1237) are separated. (3) The upper and the lower parts are rotated so that the three principal axes are aligned with three eigenvectors of the gyration tensor of the rotated points. The rotated points are smoothed by the LOWESS algorithm. (4) The upper and lower parts are rotated back to their original orientations and reconnected. The overlapping region (purple, residues 1152-1191) is averaged over the upper and lower parts. (5) The final curve is fitted by the B-spline method.

**Supplementary Figure 4.**
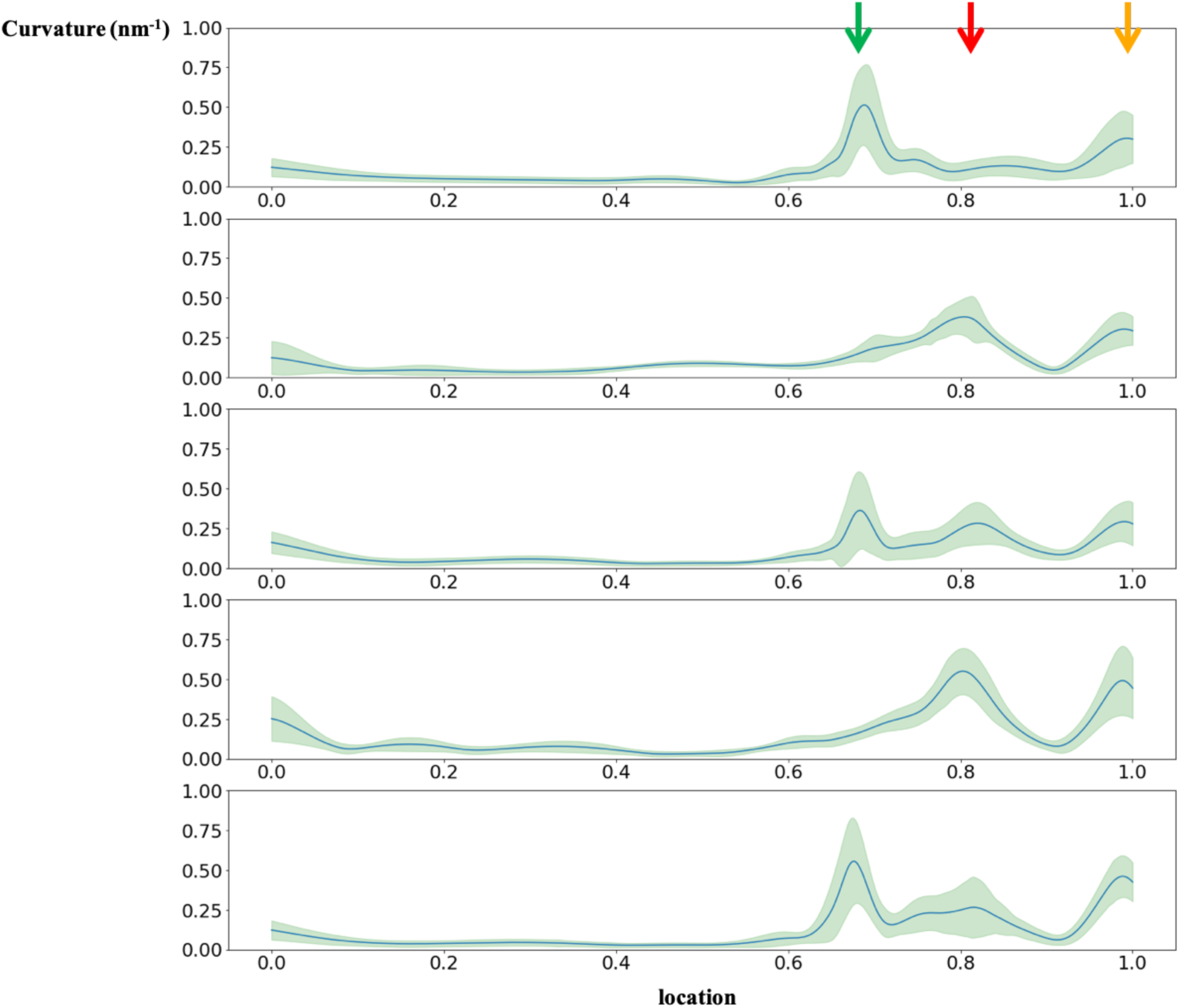
Time averaged backbone curvature versus normalized backbone arclength for five parallel runs. The three hinges (Fig. 3b, arrows) show different magnitudes of curvature in the five parallel runs. Green envelope indicates SD.

**Supplementary Figure 5.**
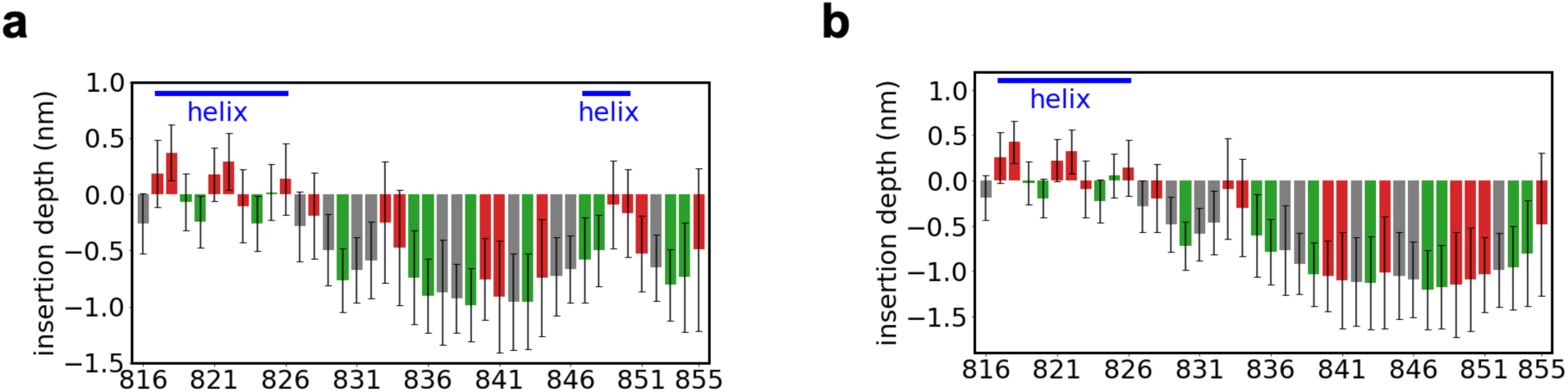
Fusion peptide residue insertion depth measured in MARTINI coarse-grained simulations. The insertion depth is defined as the vertical distance between the center of gravity (COG) of each fusion peptide residue and the COG of the PO_4_ beads in the leaflet to which the fusion peptide is bound (same definition as for the all-atom simulation). The secondary structure used is **(a)** the equilibrated secondary structure from the all-atom simulation, and **(b)** the equilibrated secondary structure but imposing the C-terminal helix to be a loop. The insertion depths are averaged over the last 78 µs. Error bars: SD over the same time frame. Bars were colored by the hydrophobicity of the corresponding residues (red: hydrophobic, grey: neutral, blue: hydrophilic), using the same color scheme as in Fig. 4b.

**Supplementary Figure 6.**
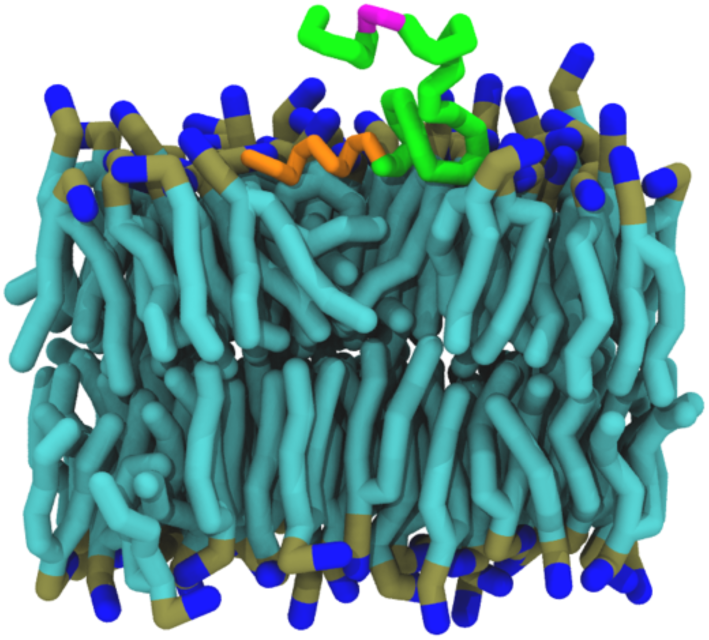
MARTINI simulation snapshot showing a transient detachment of the fusion peptide C-terminal helix. The fusion peptide C-terminal helix (purple) unanchored transiently from the membrane for ∼0.3 µs (the instance depicted here occurred after ∼52 µs of the simulation). In contrast, the N-terminal helix (orange) always remained buried in the membrane.

**Supplementary Figure 7.**
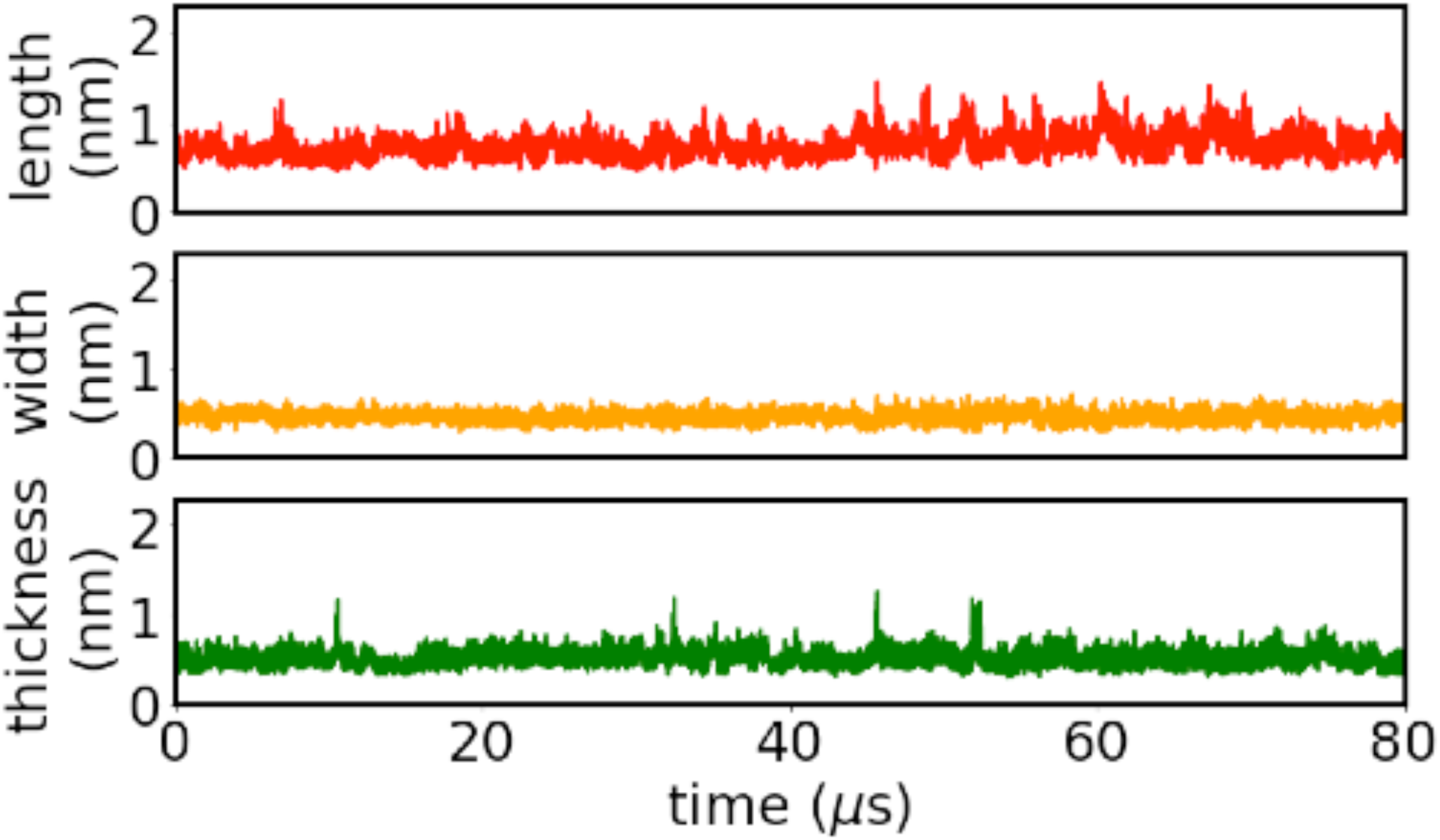
Measured length, width and thickness versus time of the fusion peptide during the MARTINI coarse-grained simulation of an equilibrated fusion peptide bound to a membrane.

**Supplementary Figure 8.**
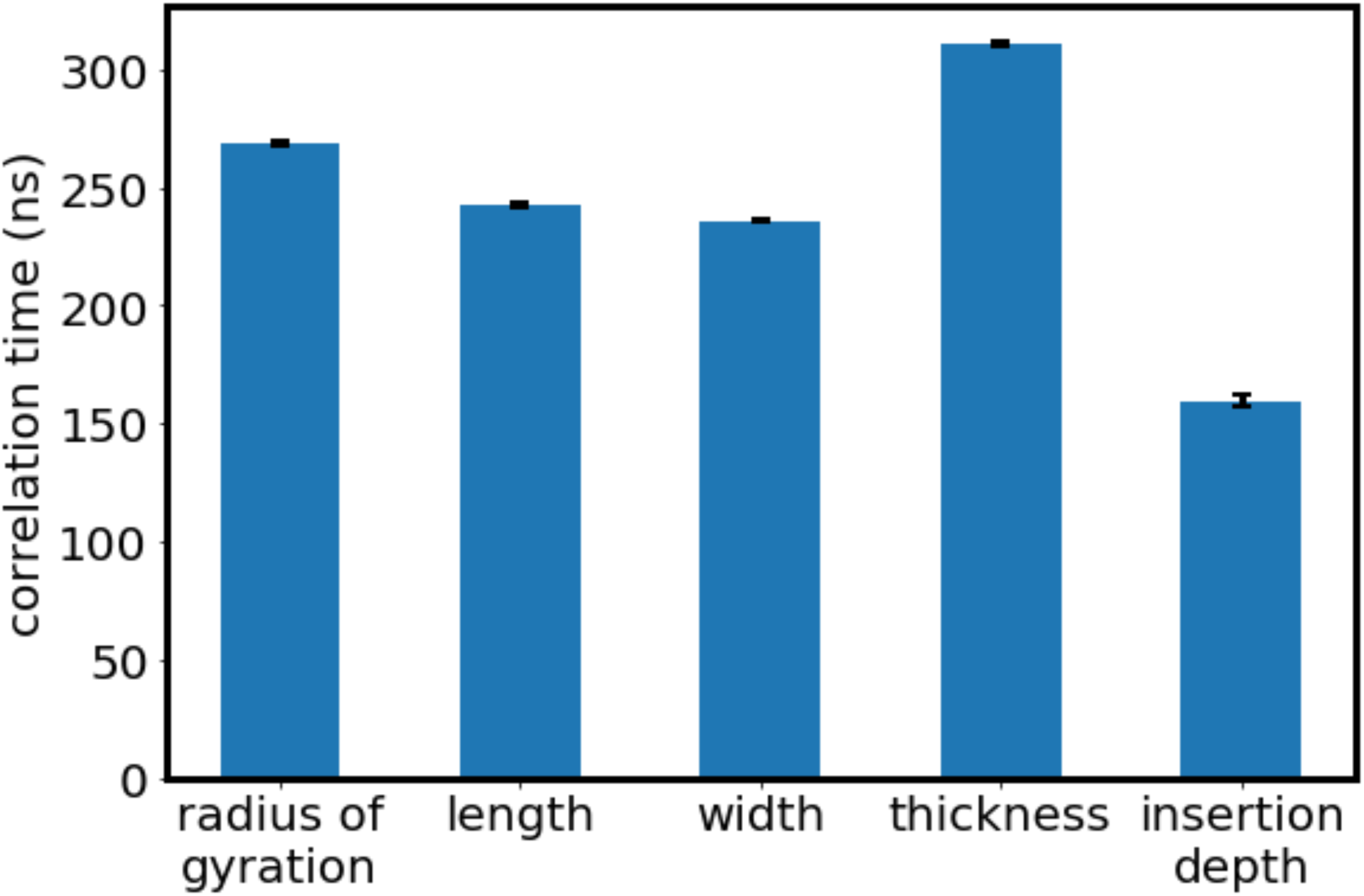
Correlation times of the fusion peptide shape properties during MARTINI coarse-grained simulation of an equilibrated fusion peptide bound to a membrane. The radius of gyration, length, width, thickness and insertion depth of the fusion peptide are calculated for the last 78 µs of the MARTINI simulation. The radius of gyration, length, width and thickness are calculated from the gyration tensor (see “Methods”). The insertion depth of the entire fusion peptide is defined as the vertical distance between the center of gravity of the fusion peptide and the center of gravity of the PO_4_ beads in the leaflet to which the FP is bound. The correlation time is calculated in the same way as in Fig. 4g. Error bars: 95% confidence interval.

**Supplementary Figure 9.**
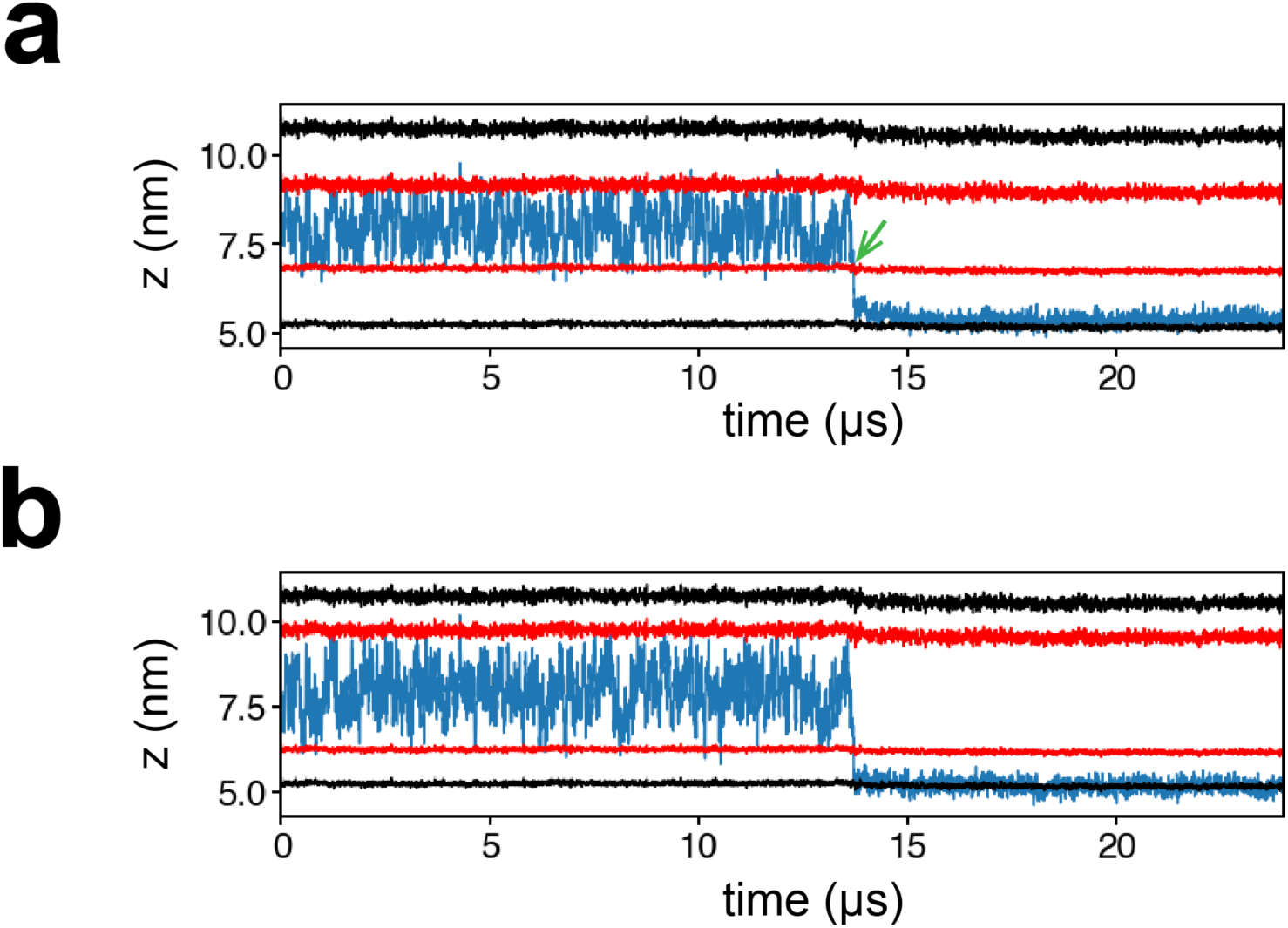
Evolution of fusion peptide vertical position during a binding assay of an isolated fusion peptide. **(a)** Evolution of the center of gravity of the fusion peptide (blue). The vertical positions of the two leaflets to which the FP is likely to bind are calculated by averaging the PO_4_ bead positions in each leaflet (black). A collision event is defined as an approach to the membrane to within *R_FP_* ∼ 1.6 nm of either leaflet (black), where *R_FP_* is the rms FP end-to-end distance. A binding event is defined to be when the FP center of gravity first has a value that positioned it below the upper membrane leaflet and above the lower membrane leaflet (green arrow). **(b)** Evolution of the center of gravity of the fusion peptide N-terminal helix (blue). The positions of the two leaflets are defined in the same way as for (a). Before the binding event, the N-terminal helix approached several times to within 1 nm of either leaflet (red).

**Supplementary Figure 10.**
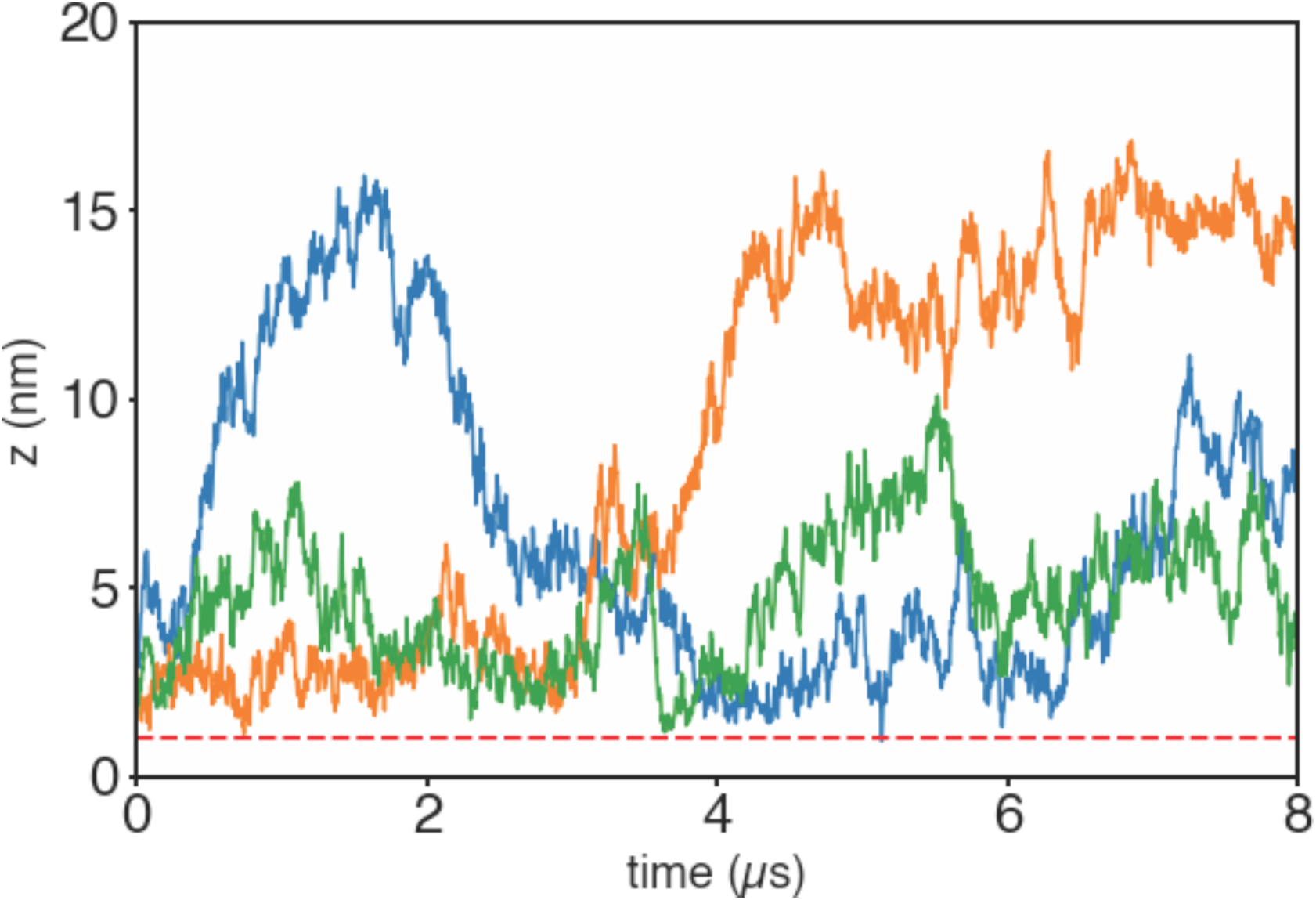
Distance of the nearest FP N-terminal helix from the membrane versus time in three runs. Here the position of the nearest FP N-terminal helix was defined to be its COG location. The membrane position was defined to be the mean location of all the PO_4_ beads in the lower leaflet of the upper membrane.

## Supplementary Movies

**Supplementary Movie 1. All-atom simulation of the SARS-CoV-2 fusion intermediate.** The color code is as for Fig. 2.

**Supplementary Movie 2. Coarse-grained MARTINI simulation of the SARS-CoV-2 fusion intermediate.** The three hinges are colored according to the code of Fig. 3c.

**Supplementary Movie 3. Simulation of Movie 2 with the fitted curve.** A fitted curve (red) to the ectodomain backbone of the fusion intermediate (blue) whose TMD is anchored in the viral envelope (green).

**Supplementary Movie 4. All-atom simulation of a membrane-bound fusion peptide (side view).** The residues are colored according to their hydrophobicity, using the color scheme of Fig. 4b.

**Supplementary Movie 5. Simulation of Movie 4, top view.**

**Supplementary Movie 6. Coarse-grained MARTINI simulation of an equilibrated fusion peptide bound to a membrane (side view).** The N-terminal helix (orange) was always buried in the membrane, but after ∼52 µs of the simulation the C-terminal helix (purple) transiently unanchored from the membrane for ∼0.3 µs.

**Supplementary Movie 7. Simulation of Movie 6, top view.**

**Supplementary Movie 8. Binding assay for an isolated fusion peptide (coarse-grained MARTINI simulation).** The fusion peptide, removed from the FI, becomes bound to the membrane after ∼1.6 µs, with the N-terminal helix (orange) providing first stable contact. The fusion peptide remains bound for the remaining ∼22.4 µs of the simulation. The binding events of Fig. 5b are snapshots from this movie.

**Supplementary Movie 9. Simulation of the fusion intermediate interacting with a target membrane, 8 μs duration.** One of the fusion peptide N-terminal helices (green spheres) repeatedly approaches the membrane to within 1 nm, but fails to bind. Fig. 6e shows a snapshot from this movie.

**Supplementary Movie 10. Binding assay of a partial fusion intermediate (coarse-grained MARTINI simulation, first 50 µs).** The partial FI consisting of the FP, CR and HR1 domains becomes bound to the membrane after ∼23 µs. The color code is as for Fig. 6b.

## References

1. Zhu, N.; Zhang, D.; Wang, W.; Li, X.; Yang, B.; Song, J.; Zhao, X.; Huang, B.; Shi, W.; Lu, R.; Niu, P.; Zhan, F.; Ma, X.; Wang, D.; Xu, W.; Wu, G.; Gao, G. F.; Tan, W., A Novel Coronavirus from Patients with Pneumonia in China, 2019. New England Journal of Medicine 2020, 382 (8), 727–733.

2. Tu, Y.-F.; Chien, C.-S.; Yarmishyn, A. A.; Lin, Y.-Y.; Luo, Y.-H.; Lin, Y.-T.; Lai, W.-Y.; Yang, D.-M.; Chou, S.-J.; Yang, Y.-P.; Wang, M.-L.; Chiou, S.-H., A Review of SARS-CoV-2 and the Ongoing Clinical Trials. International Journal of Molecular Sciences 2020, 21 (7), 2657.

3. Wrapp, D.; Wang, N.; Corbett, K. S.; Goldsmith, J. A.; Hsieh, C.-L.; Abiona, O.; Graham, B. S.; McLellan, J. S., Cryo-EM structure of the 2019-nCoV spike in the prefusion conformation. Science 2020, 367 (6483), 1260–1263.

4. Ziegler, C. G. K.; Allon, S. J.; Nyquist, S. K.; Mbano, I. M.; Miao, V. N.; Tzouanas, C. N.; Cao, Y.; Yousif, A. S.; Bals, J.; Hauser, B. M.; Feldman, J.; Muus, C.; Wadsworth, M. H.; Kazer, S. W.; Hughes, T. K.; Doran, B.; Gatter, G. J.; Vukovic, M.; Taliaferro, F.; Mead, B. E.; Guo, Z.; Wang, J. P.; Gras, D.; Plaisant, M.; Ansari, M.; Angelidis, I.; Adler, H.; Sucre, J. M. S.; Taylor, C. J.; Lin, B.; Waghray, A.; Mitsialis, V.; Dwyer, D. F.; Buchheit, K. M.; Boyce, J. A.; Barrett, N. A.; Laidlaw, T. M.; Carroll, S. L.; Colonna, L.; Tkachev, V.; Peterson, C. W.; Yu, A.; Zheng, H. B.; Gideon, H. P.; Winchell, C. G.; Lin, P. L.; Bingle, C. D.; Snapper, S. B.; Kropski, J. A.; Theis, F. J.; Schiller, H. B.; Zaragosi, L.-E.; Barbry, P.; Leslie, A.; Kiem, H.-P.; Flynn, J. L.; Fortune, S. M.; Berger, B.; Finberg, R. W.; Kean, L. S.; Garber, M.; Schmidt, A. G.; Lingwood, D.; Shalek, A. K.; Ordovas-Montanes, J.; Banovich, N.; Barbry, P.; Brazma, A.; Desai, T.; Duong, T. E.; Eickelberg, O.; Falk, C.; Farzan, M.; Glass, I.; Haniffa, M.; Horvath, P.; Hung, D.; Kaminski, N.; Krasnow, M.; Kropski, J. A.; Kuhnemund, M.; Lafyatis, R.; Lee, H.; Leroy, S.; Linnarson, S.; Lundeberg, J.; Meyer, K.; Misharin, A.; Nawijn, M.; Nikolic, M. Z.; Ordovas-Montanes, J.; Pe’er, D.; Powell, J.; Quake, S.; Rajagopal, J.; Tata, P. R.; Rawlins, E. L.; Regev, A.; Reyfman, P. A.; Rojas, M.; Rosen, O.; Saeb-Parsy, K.; Samakovlis, C.; Schiller, H.; Schultze, J. L.; Seibold, M. A.; Shalek, A. K.; Shepherd, D.; Spence, J.; Spira, A.; Sun, X.; Teichmann, S.; Theis, F.; Tsankov, A.; van den Berge, M.; von Papen, M.; Whitsett, J.; Xavier, R.; Xu, Y.; Zaragosi, L.-E.; Zhang, K., SARS-CoV-2 Receptor ACE2 Is an Interferon-Stimulated Gene in Human Airway Epithelial Cells and Is Detected in Specific Cell Subsets across Tissues. Cell 2020, 181 (5), 1016–1035.e19.

5. Hoffmann, M.; Kleine-Weber, H.; Schroeder, S.; Krüger, N.; Herrler, T.; Erichsen, S.; Schiergens, T. S.; Herrler, G.; Wu, N.-H.; Nitsche, A.; Müller, M. A.; Drosten, C.; Pöhlmann, S., SARS-CoV-2 Cell Entry Depends on ACE2 and TMPRSS2 and Is Blocked by a Clinically Proven Protease Inhibitor. Cell 2020, 181 (2), 271–280.e8.

6. Ou, X.; Liu, Y.; Lei, X.; Li, P.; Mi, D.; Ren, L.; Guo, L.; Guo, R.; Chen, T.; Hu, J.; Xiang, Z.; Mu, Z.; Chen, X.; Chen, J.; Hu, K.; Jin, Q.; Wang, J.; Qian, Z., Characterization of spike glycoprotein of SARS-CoV-2 on virus entry and its immune cross-reactivity with SARS-CoV. Nature Communications 2020, 11 (1), 1620.

7. Harrison, S. C., Viral membrane fusion. Nature Structural & Molecular Biology 2008, 15 (7), 690–698.

8. Heald-Sargent, T.; Gallagher, T., Ready, Set, Fuse! The Coronavirus Spike Protein and Acquisition of Fusion Competence. Viruses 2012, 4 (4), 557–580.

9. Cai, Y.; Zhang, J.; Xiao, T.; Peng, H.; Sterling, S. M.; Walsh, R. M.; Rawson, S.; Rits-Volloch, S.; Chen, B., Distinct conformational states of SARS-CoV-2 spike protein. Science 2020, 369 (6511), 1586–1592.

10. Jaafar, Z. A.; Kieft, J. S., Viral RNA structure-based strategies to manipulate translation. Nature Reviews Microbiology 2019, 17 (2), 110–123.

11. Walls, A. C.; Park, Y.-J.; Tortorici, M. A.; Wall, A.; McGuire, A. T.; Veesler, D., Structure, Function, and Antigenicity of the SARS-CoV-2 Spike Glycoprotein. Cell 2020, 181 (2), 281–292.e6.

12. Xia, S.; Liu, M.; Wang, C.; Xu, W.; Lan, Q.; Feng, S.; Qi, F.; Bao, L.; Du, L.; Liu, S.; Qin, C.; Sun, F.; Shi, Z.; Zhu, Y.; Jiang, S.; Lu, L., Inhibition of SARS-CoV-2 (previously 2019-nCoV) infection by a highly potent pan-coronavirus fusion inhibitor targeting its spike protein that harbors a high capacity to mediate membrane fusion. Cell Research 2020, 30 (4), 343–355.

13. Ivanovic, T.; Choi, J. L.; Whelan, S. P.; van Oijen, A. M.; Harrison, S. C., Influenza-virus membrane fusion by cooperative fold-back of stochastically induced hemagglutinin intermediates. eLife 2013, 2, e00333.

14. Ladinsky, M. S.; Gnanapragasam, P. N. P.; Yang, Z.; West, A. P.; Kay, M. S.; Bjorkman, P. J., Electron tomography visualization of HIV-1 fusion with target cells using fusion inhibitors to trap the pre-hairpin intermediate. eLife 2020, 9, e58411.

15. Benton, D. J.; Gamblin, S. J.; Rosenthal, P. B.; Skehel, J. J., Structural transitions in influenza haemagglutinin at membrane fusion pH. Nature 2020, 583 (7814), 150–153.

16. Marcink, T. C.; Wang, T.; des Georges, A.; Porotto, M.; Moscona, A., Human parainfluenza virus fusion complex glycoproteins imaged in action on authentic viral surfaces. PLOS Pathogens 2020, 16 (9), e1008883.

17. Chen, P.; Nirula, A.; Heller, B.; Gottlieb, R. L.; Boscia, J.; Morris, J.; Huhn, G.; Cardona, J.; Mocherla, B.; Stosor, V., SARS-CoV-2 neutralizing antibody LY-CoV555 in outpatients with Covid-19. New England Journal of Medicine 2021, 384 (3), 229–237.

18. Baum, A.; Ajithdoss, D.; Copin, R.; Zhou, A.; Lanza, K.; Negron, N.; Ni, M.; Wei, Y.; Mohammadi, K.; Musser, B., REGN-COV2 antibodies prevent and treat SARS-CoV-2 infection in rhesus macaques and hamsters. Science 2020, 370 (6520), 1110–1115.

19. Jackson, L. A.; Anderson, E. J.; Rouphael, N. G.; Roberts, P. C.; Makhene, M.; Coler, R. N.; McCullough, M. P.; Chappell, J. D.; Denison, M. R.; Stevens, L. J.; Pruijssers, A. J.; McDermott, A.; Flach, B.; Doria-Rose, N. A.; Corbett, K. S.; Morabito, K. M.; O’Dell, S.; Schmidt, S. D.; Swanson, P. A.; Padilla, M.; Mascola, J. R.; Neuzil, K. M.; Bennett, H.; Sun, W.; Peters, E.; Makowski, M.; Albert, J.; Cross, K.; Buchanan, W.; Pikaart-Tautges, R.; Ledgerwood, J. E.; Graham, B. S.; Beigel, J. H., An mRNA Vaccine against SARS-CoV-2 — Preliminary Report. New England Journal of Medicine 2020, 383 (20), 1920–1931.

20. Walsh, E. E.; Frenck, R. W.; Falsey, A. R.; Kitchin, N.; Absalon, J.; Gurtman, A.; Lockhart, S.; Neuzil, K.; Mulligan, M. J.; Bailey, R.; Swanson, K. A.; Li, P.; Koury, K.; Kalina, W.; Cooper, D.; Fontes-Garfias, C.; Shi, P.-Y.; Türeci, Ö.; Tompkins, K. R.; Lyke, K. E.; Raabe, V.; Dormitzer, P. R.; Jansen, K. U.; Şahin, U.; Gruber, W. C., Safety and Immunogenicity of Two RNA-Based Covid-19 Vaccine Candidates. New England Journal of Medicine 2020, 383 (25), 2439–2450.

21. de Vries, R. D.; Schmitz, K. S.; Bovier, F. T.; Predella, C.; Khao, J.; Noack, D.; Haagmans, B. L.; Herfst, S.; Stearns, K. N.; Drew-Bear, J.; Biswas, S.; Rockx, B.; McGill, G.; Dorrello, N. V.; Gellman, S. H.; Alabi, C. A.; de Swart, R. L.; Moscona, A.; Porotto, M., Intranasal fusion inhibitory lipopeptide prevents direct-contact SARS-CoV-2 transmission in ferrets. Science 2021, 371 (6536), 1379–1382.

22. Bosch, B. J.; Martina, B. E. E.; van der Zee, R.; Lepault, J.; Haijema, B. J.; Versluis, C.; Heck, A. J. R.; de Groot, R.; Osterhaus, A. D. M. E.; Rottier, P. J. M., Severe acute respiratory syndrome coronavirus (SARS-CoV) infection inhibition using spike protein heptad repeat-derived peptides. Proceedings of the National Academy of Sciences of the United States of America 2004, 101 (22), 8455–8460.

23. Marcink, T. C.; Yariv, E.; Rybkina, K.; Más, V.; Bovier, F. T.; des Georges, A.; Greninger, A. L.; Alabi, C. A.; Porotto, M.; Ben-Tal, N.; Moscona, A., Hijacking the Fusion Complex of Human Parainfluenza Virus as an Antiviral Strategy. mBio 2020, 11 (1), e03203–19.

24. LaBonte, J.; Lebbos, J.; Kirkpatrick, P., Enfuvirtide. Nature Reviews Drug Discovery 2003, 2 (5), 345.

25. Hoffmann, M.; Arora, P.; Groß, R.; Seidel, A.; Hörnich, B. F.; Hahn, A. S.; Krüger, N.; Graichen, L.; Hofmann-Winkler, H.; Kempf, A.; Winkler, M. S.; Schulz, S.; Jäck, H.-M.; Jahrsdörfer, B.; Schrezenmeier, H.; Müller, M.; Kleger, A.; Münch, J.; Pöhlmann, S., SARS-CoV-2 variants B.1.351 and P.1 escape from neutralizing antibodies. Cell 2021.

26. Tang, T.; Bidon, M.; Jaimes, J. A.; Whittaker, G. R.; Daniel, S., Coronavirus membrane fusion mechanism offers as a potential target for antiviral development. Antiviral Research 2020, 104792.

27. Casalino, L.; Gaieb, Z.; Goldsmith, J. A.; Hjorth, C. K.; Dommer, A. C.; Harbison, A. M.; Fogarty, C. A.; Barros, E. P.; Taylor, B. C.; McLellan, J. S.; Fadda, E.; Amaro, R. E., Beyond Shielding: The Roles of Glycans in the SARS-CoV-2 Spike Protein. ACS Central Science 2020, 6 (10), 1722–1734.

28. Turoňová, B.; Sikora, M.; Schürmann, C.; Hagen, W. J. H.; Welsch, S.; Blanc, F. E. C.; von Bülow, S.; Gecht, M.; Bagola, K.; Hörner, C.; van Zandbergen, G.; Landry, J.; de Azevedo, N. T. D.; Mosalaganti, S.; Schwarz, A.; Covino, R.; Mühlebach, M. D.; Hummer, G.; Krijnse Locker, J.; Beck, M., In situ structural analysis of SARS-CoV-2 spike reveals flexibility mediated by three hinges. Science 2020, 370 (6513), 203–208.

29. Ali, A.; Vijayan, R., Dynamics of the ACE2–SARS-CoV-2/SARS-CoV spike protein interface reveal unique mechanisms. Scientific Reports 2020, 10 (1), 14214.

30. Khelashvili, G.; Plante, A.; Doktorova, M.; Weinstein, H., Ca^2+^-dependent mechanism of membrane insertion and destabilization by the SARS-CoV-2 fusion peptide. Biophysical Journal 2021, 120 (6), 1105–1119.

31. Gorgun, D.; Lihan, M.; Kapoor, K.; Tajkhorshid, E., Binding mode of SARS-CoV-2 fusion peptide to human cellular membrane. Biophysical Journal 2021.

32. Borkotoky, S.; Dey, D.; Banerjee, M., Computational Insight Into the Mechanism of SARS-CoV-2 Membrane Fusion. Journal of Chemical Information and Modeling 2021, 61 (1), 423–431.

33. Yu, A.; Pak, A. J.; He, P.; Monje-Galvan, V.; Casalino, L.; Gaieb, Z.; Dommer, A. C.; Amaro, R. E.; Voth, G. A., A multiscale coarse-grained model of the SARS-CoV-2 virion. Biophysical Journal 2021, 120 (6), 1097–1104.

34. Barfoot, S.; Poger, D.; Mark, A. E., Understanding the Activated Form of a Class-I Fusion Protein: Modeling the Interaction of the Ebola Virus Glycoprotein 2 with a Lipid Bilayer. Biochemistry 2020, 59 (41), 4051–4058.

35. Lin, X.; Noel, J. K.; Wang, Q.; Ma, J.; Onuchic, J. N., Atomistic simulations indicate the functional loop-to-coiled-coil transition in influenza hemagglutinin is not downhill. Proceedings of the National Academy of Sciences 2018, 115 (34), E7905–E7913.

36. Lin, M.; Da, L.-T., Refolding Dynamics of gp41 from Pre-fusion to Pre-hairpin States during HIV-1 Entry. Journal of Chemical Information and Modeling 2020, 60 (1), 162–174.

37. Carr, C. M.; Kim, P. S., A spring-loaded mechanism for the conformational change of influenza hemagglutinin. Cell 1993, 73 (4), 823–832.

38. Benton, D. J.; Wrobel, A. G.; Xu, P.; Roustan, C.; Martin, S. R.; Rosenthal, P. B.; Skehel, J. J.; Gamblin, S. J., Receptor binding and priming of the spike protein of SARS-CoV-2 for membrane fusion. Nature 2020, 588 (7837), 327–330.

39. Hakansson-McReynolds, S.; Jiang, S.; Rong, L.; Caffrey, M., Solution Structure of the Severe Acute Respiratory Syndrome-Coronavirus Heptad Repeat 2 Domain in the Prefusion State *. Journal of Biological Chemistry 2006, 281 (17), 11965–11971.

40. Xu, D.; Zhang, Y., Ab initio protein structure assembly using continuous structure fragments and optimized knowledge-based force field. Proteins: Structure, Function, and Bioinformatics 2012, 80 (7), 1715–1735.

41. Ke, Z.; Oton, J.; Qu, K.; Cortese, M.; Zila, V.; McKeane, L.; Nakane, T.; Zivanov, J.; Neufeldt, C. J.; Cerikan, B.; Lu, J. M.; Peukes, J.; Xiong, X.; Kräusslich, H.-G.; Scheres, S. H. W.; Bartenschlager, R.; Briggs, J. A. G., Structures and distributions of SARS-CoV-2 spike proteins on intact virions. Nature 2020, 588 (7838), 498–502.

42. Lai, A. L.; Freed, J. H., SARS-CoV-2 Fusion Peptide has a Greater Membrane Perturbating Effect than SARS-CoV with Highly Specific Dependence on Ca2+. Journal of Molecular Biology 2021, 433 (10), 166946.

43. O’Shaughnessy, B.; Sawhney, U., Polymer Reaction Kinetics at Interfaces. Physical Review Letters 1996, 76 (18), 3444–3447.

44. Outlaw, V. K.; Bovier, F. T.; Mears, M. C.; Cajimat, M. N.; Zhu, Y.; Lin, M. J.; Addetia, A.; Lieberman, N. A. P.; Peddu, V.; Xie, X.; Shi, P.-Y.; Greninger, A. L.; Gellman, S. H.; Bente, D. A.; Moscona, A.; Porotto, M., Inhibition of Coronavirus Entry *In Vitro* and *Ex Vivo* by a Lipid-Conjugated Peptide Derived from the SARS-CoV-2 Spike Glycoprotein HRC Domain. mBio 2020, 11 (5), e01935–20.

45. Baylon, J. L.; Tajkhorshid, E., Capturing Spontaneous Membrane Insertion of the Influenza Virus Hemagglutinin Fusion Peptide. The Journal of Physical Chemistry B 2015, 119 (25), 7882–7893.

46. Calder, L. J.; Rosenthal, P. B., Cryomicroscopy provides structural snapshots of influenza virus membrane fusion. Nature Structural & Molecular Biology 2016, 23 (9), 853–858.

47. Benton, D. J.; Nans, A.; Calder, L. J.; Turner, J.; Neu, U.; Lin, Y. P.; Ketelaars, E.; Kallewaard, N. L.; Corti, D.; Lanzavecchia, A.; Gamblin, S. J.; Rosenthal, P. B.; Skehel, J. J., Influenza hemagglutinin membrane anchor. Proceedings of the National Academy of Sciences 2018, 115 (40), 10112–10117.

48. Lu, L.; Liu, Q.; Zhu, Y.; Chan, K.-H.; Qin, L.; Li, Y.; Wang, Q.; Chan, J. F.-W.; Du, L.; Yu, F.; Ma, C.; Ye, S.; Yuen, K.-Y.; Zhang, R.; Jiang, S., Structure-based discovery of Middle East respiratory syndrome coronavirus fusion inhibitor. Nature Communications 2014, 5 (1), 3067.

49. Wang, C.; van Haperen, R.; Gutiérrez-Álvarez, J.; Li, W.; Okba, N. M. A.; Albulescu, I.; Widjaja, I.; van Dieren, B.; Fernandez-Delgado, R.; Sola, I.; Hurdiss, D. L.; Daramola, O.; Grosveld, F.; van Kuppeveld, F. J. M.; Haagmans, B. L.; Enjuanes, L.; Drabek, D.; Bosch, B.-J., A conserved immunogenic and vulnerable site on the coronavirus spike protein delineated by cross-reactive monoclonal antibodies. Nature Communications 2021, 12 (1), 1715.

50. Webb, B.; Sali, A., Comparative Protein Structure Modeling Using MODELLER. Curr Protoc Bioinformatics 2016, 54 (1), 5.6.1-5.6.37.

51. Bekker, H.; Berendsen, H.; Dijkstra, E.; Achterop, S.; Van Drunen, R.; Van der Spoel, D.; Sijbers, A.; Keegstra, H.; Reitsma, B.; Renardus, M. In *Gromacs: A parallel computer for molecular dynamics simulations*, Physics computing, World Scientific Singapore: 1993; pp 252–256.

52. Berendsen, H. J.; van der Spoel, D.; van Drunen, R. J. C. p. c., GROMACS: a message-passing parallel molecular dynamics implementation. 1995, 91 (1-3), 43–56.

53. Jo, S.; Kim, T.; Iyer, V. G.; Im, W., CHARMM-GUI: A web-based graphical user interface for CHARMM. Journal of Computational Chemistry 2008, 29 (11), 1859–1865.

54. Best, R. B.; Zhu, X.; Shim, J.; Lopes, P. E.; Mittal, J.; Feig, M.; MacKerell Jr, A. D. J. J. o. c. t.; computation, Optimization of the additive CHARMM all-atom protein force field targeting improved sampling of the backbone ϕ, ψ and side-chain χ1 and χ2 dihedral angles. 2012, 8 (9), 3257–3273.

55. Klauda, J. B.; Venable, R. M.; Freites, J. A.; O’Connor, J. W.; Tobias, D. J.; Mondragon-Ramirez, C.; Vorobyov, I.; MacKerell Jr, A. D.; Pastor, R. W. J. T. j. o. p. c. B., Update of the CHARMM all-atom additive force field for lipids: validation on six lipid types. 2010, 114 (23), 7830–7843.

56. Kabsch, W.; Sander, C., Dictionary of protein secondary structure: pattern recognition of hydrogen-bonded and geometrical features. Biopolymers 1983, 22 (12), 2577–637.

57. Joosten, R. P.; Te Beek, T. A.; Krieger, E.; Hekkelman, M. L.; Hooft, R. W.; Schneider, R.; Sander, C.; Vriend, G. J. N. a. r., A series of PDB related databases for everyday needs. 2010, 39 (suppl_1), D411–D419.

58. Wassenaar, T. A.; Pluhackova, K.; Böckmann, R. A.; Marrink, S. J.; Tieleman, D. P. J. J. o. c. t.; computation, Going backward: a flexible geometric approach to reverse transformation from coarse grained to atomistic models. 2014, 10 (2), 676–690.

59. Marrink, S. J.; de Vries, A. H.; Mark, A. E., Coarse Grained Model for Semiquantitative Lipid Simulations. The Journal of Physical Chemistry B 2004, 108 (2), 750–760.

60. de Jong, D. H.; Singh, G.; Bennett, W. D.; Arnarez, C.; Wassenaar, T. A.; Schafer, L. V.; Periole, X.; Tieleman, D. P.; Marrink, S. J. J. J. o. c. t.; computation, Improved parameters for the martini coarse-grained protein force field. 2013, 9 (1), 687–697.

61. Marrink, S. J.; Risselada, H. J.; Yefimov, S.; Tieleman, D. P.; De Vries, A. H. J. T. j. o. p. c. B., The MARTINI force field: coarse grained model for biomolecular simulations. 2007, 111 (27), 7812–7824.

62. Wassenaar, T. A.; Ingólfsson, H. I.; Böckmann, R. A.; Tieleman, D. P.; Marrink, S. J. J. J. o. c. t.; computation, Computational lipidomics with insane: a versatile tool for generating custom membranes for molecular simulations. 2015, 11 (5), 2144–2155.

## Supplementary References

1 Jo, S., Kim, T., Iyer, V. G. & Im, W. J. J. o. c. c. CHARMM-GUI: a web-based graphical user interface for CHARMM. 29, 1859–1865 (2008).

2 Jorgensen, W. L., Chandrasekhar, J., Madura, J. D., Impey, R. W. & Klein, M. L. J. T. J. o. c. p. Comparison of simple potential functions for simulating liquid water. 79, 926–935 (1983).

3 Hoover, W. G. J. P. r. A. Canonical dynamics: Equilibrium phase-space distributions. 31, 1695 (1985).

4 Nosé, S. J. T. J. o. c. p. A unified formulation of the constant temperature molecular dynamics methods. 81, 511–519 (1984).

5 Parrinello, M. & Rahman, A. J. J. o. A. p. Polymorphic transitions in single crystals: A new molecular dynamics method. 52, 7182–7190 (1981).

6 Bekker, H. et al. in Physics computing. 252–256 (World Scientific Singapore).

7 Berendsen, H. J., van der Spoel, D. & van Drunen, R. J. C. p. c. GROMACS: a message-passing parallel molecular dynamics implementation. 91, 43–56 (1995).

8 Kabsch, W. & Sander, C. J. B. O. R. o. B. Dictionary of protein secondary structure: pattern recognition of hydrogen-bonded and geometrical features. 22, 2577–2637 (1983).

9 Joosten, R. P., et al. A series of PDB related databases for everyday needs. 39, D411–D419 (2010).

10 Cai, Y. et al. Distinct conformational states of SARS-CoV-2 spike protein. 369, 1586–1592 (2020).

11 Best, R. B. et al. Optimization of the additive CHARMM all-atom protein force field targeting improved sampling of the backbone ϕ, ψ and side-chain χ1 and χ2 dihedral angles. 8, 3257–3273 (2012).

12 Klauda, J. B. et al. Update of the CHARMM all-atom additive force field for lipids: validation on six lipid types. 114, 7830–7843 (2010).

13 de Jong, D. H. et al. Improved parameters for the martini coarse-grained protein force field. 9, 687–697 (2013).

14 Marrink, S. J., Risselada, H. J., Yefimov, S., Tieleman, D. P. & De Vries, A. H. J. T. j. o. p. c. B. The MARTINI force field: coarse grained model for biomolecular simulations. 111, 7812–7824 (2007).

15 Wassenaar, T. A. et al. Computational lipidomics with insane: a versatile tool for generating custom membranes for molecular simulations. 11, 2144–2155 (2015).

16 Bussi, G., Donadio, D. & Parrinello, M. J. T. J. o. c. p. Canonical sampling through velocity rescaling. 126, 014101 (2007).

17 Berendsen, H. J., Postma, J. v., van Gunsteren, W. F., DiNola, A. & Haak, J. R. J. T. J. o. c. p. Molecular dynamics with coupling to an external bath. 81, 3684–3690 (1984).

18 McGibbon, R. T. et al. MDTraj: a modern open library for the analysis of molecular dynamics trajectories. 109, 1528–1532 (2015).

19 Wassenaar, T. A. et al. Going backward: a flexible geometric approach to reverse transformation from coarse grained to atomistic models. 10, 676–690 (2014).

